# A tunable dual-input system for ‘on-demand’ dynamic gene expression regulation

**DOI:** 10.1101/404699

**Authors:** Elisa Pedone, Dan L. Rocca, Lorena Postiglione, Francesco Aulicino, Sandra Montes-Olivas, Diego di Bernardo, Maria Pia Cosma, Lucia Marucci

## Abstract

Cellular systems have evolved numerous mechanisms to finely control signalling pathway activation and properly respond to changing environmental stimuli. This is underpinned by dynamic spatiotemporal patterns of gene expression. Indeed, in addition to gene transcription and translation regulation, modulation of protein levels, dynamics and localization are also essential checkpoints that govern cell functions. The introduction of tetracycline-inducible promoters has allowed gene expression control using orthogonal small molecules, facilitating rapid and reversible manipulation to study gene function in biological systems. However, differing protein stabilities means this solely transcriptional regulation is insufficient to allow precise ON-OFF dynamics, thus hindering generation of temporal profiles of protein levels seen *in vivo*. We developed an improved Tet-On based system augmented with conditional destabilising elements at the post-translational level that permits simultaneous control of gene expression and protein stability. Integrating these properties to control expression of a fluorescent protein in mouse Embryonic Stem Cells (mESCs), we found that adding protein stability control allows faster response times to changes in small molecules, fully tunable and enhanced dynamic range, and vastly improved microfluidic-based *in-silico* feedback control of gene expression. Finally, we highlight the effectiveness of our dual-input system to finely modulate levels of signalling pathway components in stem cells.

## Introduction

A number of perturbation approaches has been developed to study gene function in biological systems, and for gene therapy applications. It has become increasingly clear that gene expression patterns *in vivo* are fast and highly dynamic processes, encoding important time-dependent information that underlies many aspects of cellular behaviour^1^. Thus, precise temporal control, as well as reversible manipulation of exogenous gene expression is fundamental for interrogating cellular functions^2^. The ability to turn the expression of transgenes ON and OFF, or to finely modulate their expression levels, could greatly improve safety in gene therapy strategies by reducing unwanted off-targets and side effects^3^.

Temporal control of gene expression can be achieved by transcriptional regulation *via* inducible promoters^4-7^.The most widely used system for transcriptional regulation is the Tet system, which consists of two elements: the tetracycline transcriptional activator (tTA) and the DNA operator sequence (tetO)^8-12^. The tTA is a fusion protein of the herpes simplex virus VP16 activation domain and of the *Escherichia coli* Tet repressor protein (TetR)^9^. The presence of tetracycline or its derivative doxycycline prevents the interaction of the tTA to the tetO, blocking gene expression (Tet-Off system). The reverse-tTA (rtTA) is a tTA variant allowing gene expression activation in presence of an inducer; the resulting Tet-On system is generally preferred when rapid and dynamic gene induction is required^4,13^. A major limitation of inducible promoters is the significant time delay in switching proteins OFF and ON when using Tet-On and Tet-off systems, respectively^14^, diminishing the possibility of using these approaches to generate dynamic patterns of gene expression that faithfully recapitulate those observed natively^1^. Slow kinetic responses are also common to other techniques targeting precursor DNA or mRNA molecules (e.g. RNA interference^15^), likely due to significantly different rates of innate protein degradation^15^.

Recently, an alternative approach, relying on conditional protein destabilization to modulate turnover by the cellular degradation machinery, has been harnessed to probe complex biological functions. Engineered mutants of FKBP12 that are rapidly and constitutively degraded in mammalian cells can directly confer protein destabilization to the protein they are fused with. Addition of synthetic ligands, that bind the Destabilising Domain (DD) of FKBP12, prevent degradation and so can be used to alter levels of the fused-protein of interest^16^. The plant derived AID (auxin-inducible degradation) system is also used to get fast and efficient proteasomal degradation of AID-tagged proteins in response to auxin hormones^17^. However, rates of AID-mediated protein degradation/recovery strongly depend on auxin uptake and metabolism, as well as on the abundance of SCF complex components, which might vary among different biological systems^18,19^. Therefore, AID is more suitable for protein knockout experiments, whereas the DD-based tool is preferred when fine temporal control of fused-protein levels is needed.

Whilst significantly enhancing the switch-off kinetics as compared to Tet-On regulation, conditional protein regulation systems do not allow independent control of both transcription and translation, which would be highly desirable when studying the correlation between protein and cognate mRNA levels under different spatial and temporal scales^20^.

To overcome these limitations, we engineered a novel, fully tuneable dual-input system, which allows orthogonal and conditional control at both transcriptional and post-translational levels of a gene of interest. Specifically, we combined a third generation Tetracycline-Inducible System (Tet-ON 3G)^12,21^ for inducible and reversible transcriptional regulation with a module incorporating an improved DD from ecDHFR^22^ for targeted protein degradation. We demonstrated that our system permits far greater control of both protein dynamics and expression dynamic range across different culture platforms, including microfluidics used for *in silico* feedback control. Moreover, we developed an ordinary differential equation model, which captures the enhanced dynamic response to inducers. The efficacy of conditional, dual regulation inherent in our system is exemplified by the ability to incorporate different genes of interest, such as fluorescent proteins, as well as the Wnt signalling effector, β-catenin, in a complex cellular chassis (mouse embryonic stem cells), paving the way for dynamically controlling mammalian cell behaviour and fate.

## Results

### Dual-input system for orthogonal regulation of transcriptional and post-translational gene expression

We engineered a mouse Embryonic Stem Cell (mESC) line to stably express the reverse tetracycline transcriptional activator construct (rtTA) and a stable mCherry (henceforth EF1a-rtTA_TRE3G-mCherry; Fig. 1a), or conditionally destabilized DDmCherry (henceforth EF1a-rtTA_TRE3G-DDmCherry; Fig. 1c) under the control of a TRE3G promoter, which transcribes the gene of interest only in presence of the tetracycline analogue doxycycline^12^ (Doxy; Fig. 1a, c). Post-translational control is achieved by applying the small molecule trimethoprim (TMP), which stabilizes the destabilizing domain (DD)-fused protein in a dose-dependent manner^22^. The two constructs allow for comparison of the standard Tet-on with the dual-input Tet-On/DD system we developed.

**Figure 1.**
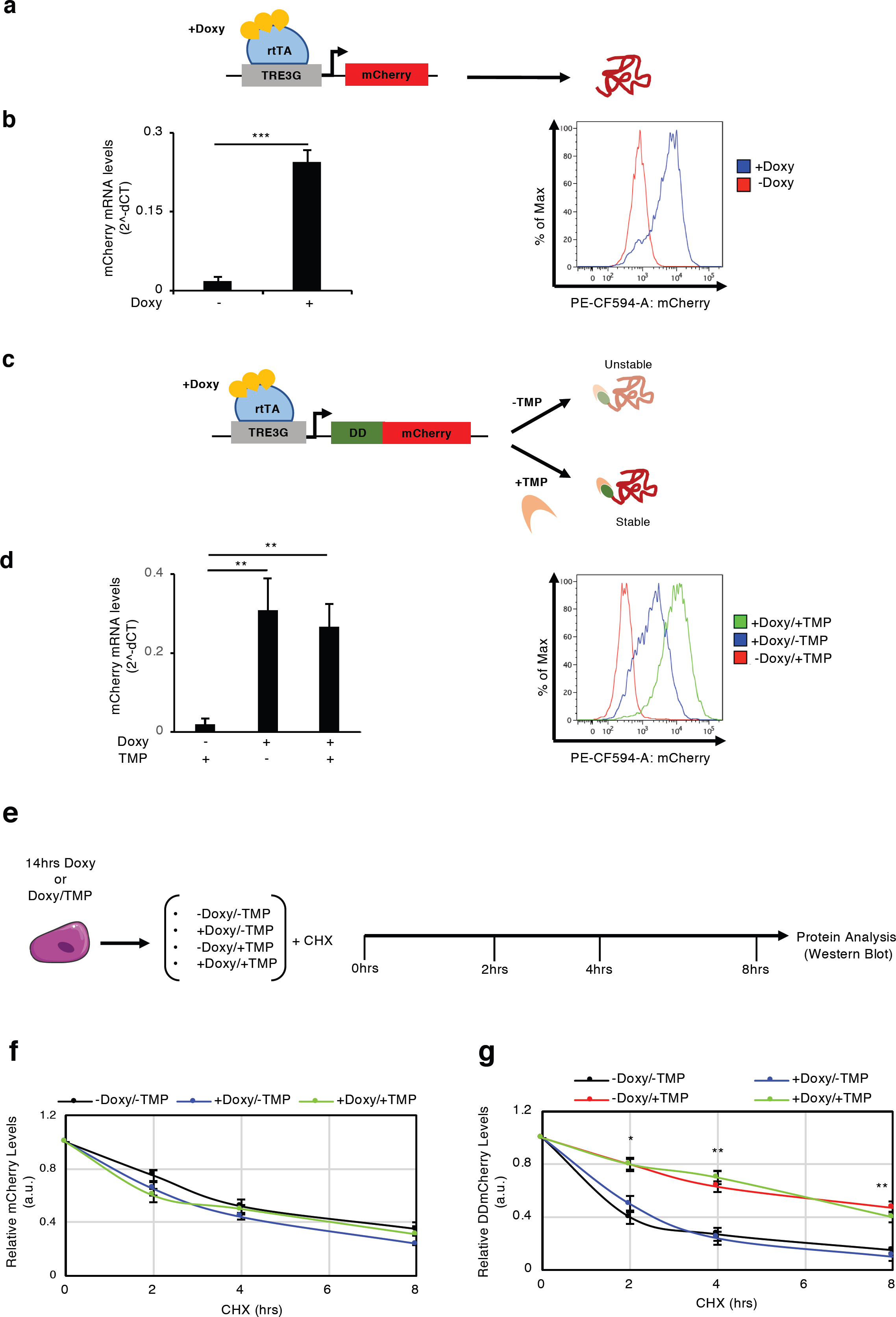
Dual-input regulation of exogenous protein expression. **(a-d)** Dual-input regulation system consisting of a reverse transactivator (rtTA) and stable **(a)** or conditionally destabilised **(c)** mCherry fluorescent protein. mRNA **(b, d, left panels)** and protein levels **(b, d, right panels)** measured in EF1a-rtTA_TRE3G-mCherry **(b)** and EF1a-rtTA_TRE3G-DDmCherry **(d)** mESCs treated for 24hrs with Doxy (1000ng/mL) or Doxy/TMP (1000ng/mL and 100nM, respectively). **(e)** Experimental scheme of protein half-life measurement experiments. Following 14hrs of treatment with Doxy (1000ng/mL) or Doxy/TMP (1000ng/mL and 100nM, respectively), EF1a-rtTA_TRE3G-mCherry and EF1a-rtTA_TRE3G-DDmCherry mESCs were cultured in presence of the protein synthesis inhibitor cycloheximide (CHX, 25μg/mL) and combination of Doxy (1000ng/mL), TMP (100nM) and Doxy/TMP. Protein half-life was measured by SDS-PAGE and western-blotting after the indicated times of treatment. **(f, g)** Western-blot densitometric quantification of EF1a-rtTA_TRE3G-mCherry **(f)** and EF1a-rtTA_TRE3G-DDmCherry **(g)** mESCs. Data are means ± SEM (n = 2, **b** and **d**; n=3, **f** and **g**). p > 0.1, *p < 0.05, **p < 0.01,***p < 0.0001.

We exposed EF1a-rtTA_TRE3G-mCherry and EF1a-rtTA_TRE3G-DDmCherry mESCs to saturating concentrations of Doxy (1000ng/mL) and TMP (100nM) for 24hrs and checked for mCherry mRNA (Fig. 1b, d, left panels) and protein (Fig. 1b, d, right panels) levels. In absence of Doxy, both transcription and protein expression were undetectable, demonstrating no promoter leakiness (Fig. 1b, d; -Doxy and - Doxy/+TMP samples, respectively), whereas mCherry was robustly activated at both mRNA and protein levels following Doxy and TMP administration (Fig. 1b, d; +Doxy and +Doxy/±TMP, respectively). Notably, Doxy and TMP selectively control transcription and protein stability, respectively: when TMP is combined with Doxy, mCherry mRNA is unchanged, (Fig. 1d; left panel), whereas a clear increase in protein stability is observed (Fig. 1d; right panel).

To quantitatively estimate the effect of inducers on protein stability, we measured protein half-life in both EF1a-rtTA_TRE3G-mCherry and EF1a-rtTA_TRE3G-DDmCherry mESCs. Cells were cultured in presence of Doxy and Doxy/TMP for 14hrs, and then plated in presence of the protein synthesis inhibitor cycloheximide with varying combinations of Doxy and TMP for 8hrs; mCherry was measured by Western Blot over the time-course (Fig. 1e). In EF1a-rtTA_TRE3G-mCherry mESCs, untagged mCherry showed no response to TMP, and the half-life (around 4hrs) was similar across conditions (Fig. 1f; Supplementary Fig.1a). Of note, mCherry mRNA decreased following Doxy withdrawal (Supplementary Fig. 1c), confirming no effect of Doxy or TMP on protein stability in this system. In contrast, the half-life of the conditionally destabilised mCherry increased approximately three fold in presence of TMP (Fig. 1g; Supplementary Fig.1b; +Doxy/+TMP and -Doxy/+TMP samples). In addition, mRNA levels decreased only when Doxy was removed (Supplementary Fig. 1d), further demonstrating the specific and independent effect of the two inducers.

Consistently, we found that MG132 blockade of the proteasome^23^ enhanced DDmCherry stability, suggesting that TMP acts by preventing proteasomal degradation of DD-fused proteins only^22^ (Supplementary Fig. 1e). Indeed, both MG132 and TMP treatment increased DDmCherry abundance, whereas levels of the native β-catenin, whose degradation is proteasome mediated^24^, increased only when proteasomal activity was inhibited (Supplementary Fig. 1e). Analysing the polyubiquitination status of DDmCherry, following inducer removal and chasing with or without TMP for 12hrs, indicated that TMP likely limits addition of ubiquitin chains or promotes de-ubiquitination of DD-tagged proteins, ultimately preventing proteasomal degradation (Supplementary Fig. 1f).

Altogether, these data show that our system allows robust control of both gene transcription and protein stability, with undetectable leakiness and specific response to inducers.

### Dual-input system tunability and dynamic response

To gauge the tunability and sensitivity of our inducible system, as well as its suitability for dynamic modulation of a gene of interest, we performed steady-state titration experiments with both drugs, and measured the switch-off dynamics of EF1a-rtTA_TRE3G-mCherry and EF1a-rtTA_TRE3G-DDmCherry mESCs in flow-cytometry time-courses.

EF1a-rtTA_TRE3G-mCherry mESCs incubated with 6 different concentrations of Doxy for 24hrs showed a robust response to increasing amount of inducer, with saturation reached at Doxy 100ng/ml (Fig. 2a, dots; Supplementary Fig. 2a); EF1a-rtTA_TRE3G-DDmCherry mESCs showed similar steady-state response, when kept with saturating concentration of TMP (10μM) and varying Doxy in the same range (Fig. 2b, dots; Supplementary Fig. 2b). Also, TMP showed a robust dose-dependent effect, when varied while keeping Doxy at maximal concentration (Fig. 2c, dots; Supplementary Fig. 2b).

**Figure 2.**
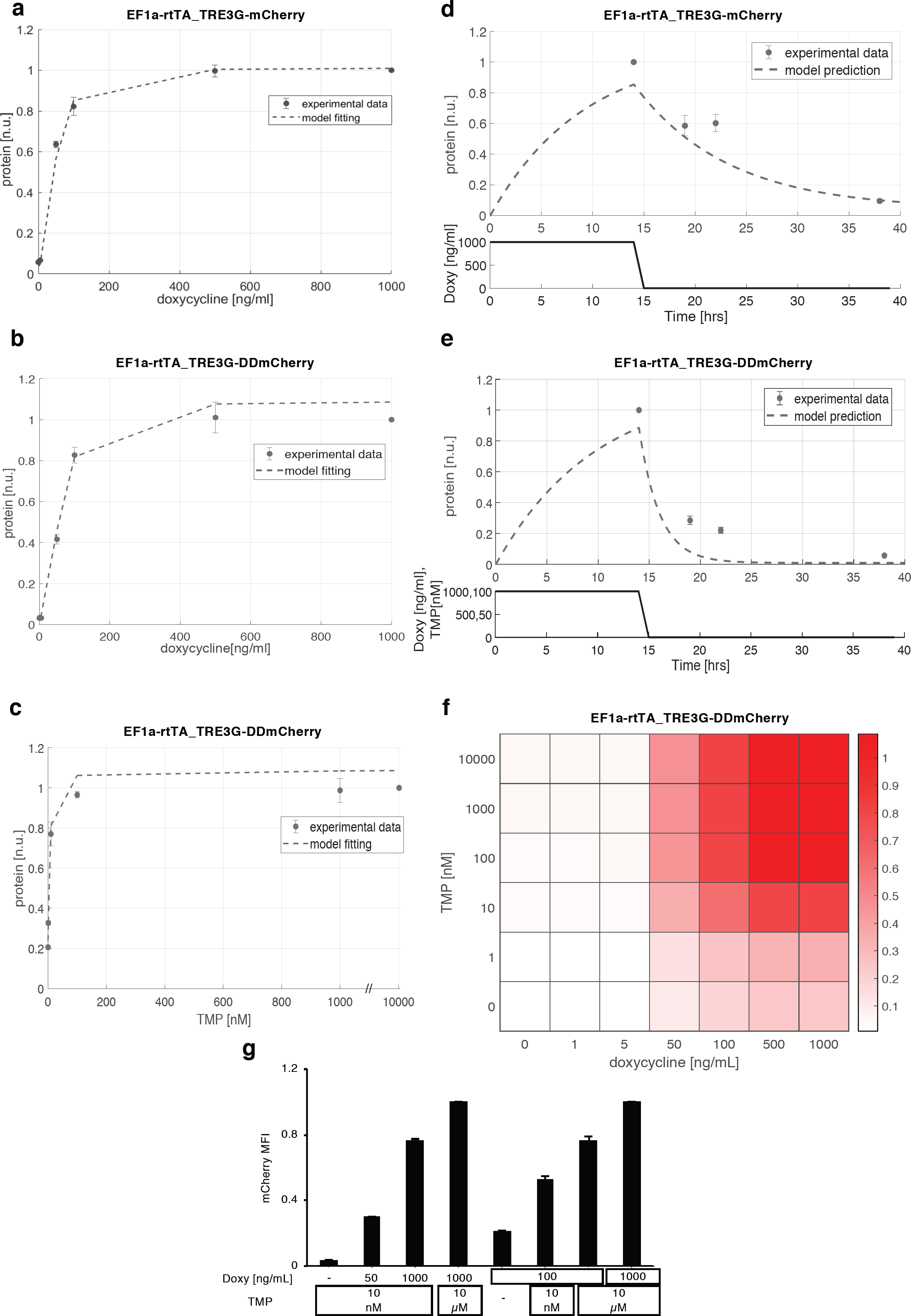
Steady-state/dynamic response and dynamic range of the dual-input system. **(a)** mCherry protein levels in EF1a-rtTA_TRE3G-mCherry mESCs following 24hrs of treatment with increasing concentration of Doxy (0; 1; 5; 50; 100; 500; 1000ng/mL). **(b, c)** mCherry protein levels measured in EF1a-rtTA_TRE3G-DDmCherry following 24hrs of treatment with increasing concentration of Doxy (0; 1; 5; 50; 100; 500; 1000ng/mL) and saturated TMP (10μM) **(b),** or increasing concentration of TMP (0; 1; 10; 100nM; 1; 10μM) and saturated Doxy (1000ng/mL) **(c)**. **(d, e)** Dynamic response of EF1a-rtTA_TRE3G-mCherry **(d)** and EF1a-rtTA_TRE3G-DDmCherry **(e)** mESCs to Doxy (1000ng/mL) or Doxy/TMP (1000ng/mL and 100nM, respectively) in a time-course experiment of 38hrs, in which inducers were washed out after incubation in the first 14hrs. The dots are experimental data of MFI (Supplementary Fig. 2a-d) measured by flow cytometry and normalised over the maximum activation point. Dashed lines represent model fitting of steady-state data **(a-c)** or model prediction of switch-off dynamics **(d, e)**. **(f)** Simulated steady-state of EF1a-rtTA_TRE3G-DDmCherry mESCs upon combined treatment with Doxy and TMP at various concentrations. mCherry values are shown as heatmaps of scaled values across the entire dynamic range of expression levels. **(g)** Experimental validation of simulations in **(f)**, measuring EF1a-rtTA_TRE3G-DDmCherry mESC steady-state following 24hrs induction with constant concentration of Doxy (100ng/mL) and varying TMP (0; 10nM; 10μM), or constant TMP (10nM) and varying Doxy (0; 50; 1000ng/mL). Maximum concentrations of Doxy and TMP (1000mg/mL and 10μM, respectively) are used as control. Data are means ± SD (n=3, **a-f**); ± SEM (n=3, **g**).

To complement these experiments, we developed a mathematical model to capture the behaviour of the dual-input system: we relied on ordinary differential equations (ODEs), commonly used to model interactions among genes and other relevant processes, such as mRNA/protein degradation, basal promoter activity and mRNA translation^25,26^. The mathematical models for the EF1a-rtTA_TRE3G-mCherry and EF1a-rtTA_TRE3G-DDmCherry systems are based on sets of 4 ODEs, describing transactivator and fluorescent gene mRNA and corresponding protein concentrations, as the result of production and degradation terms (Supplementary Information). The TET system was modelled, as previously proposed, using Hill kinetics to represent the effect of the inducer on trascription^27^. Given the observed saturating response to TMP (Fig. 2c, dots and ^22^), a Hill function, dependent on TMP, was incorporated in the DDmCherry protein degradation term (Supplementary Information). To estimate parameters (reported in Table 1, Supplementary Information), the two ODE models were fitted to experimental data and showed good agreement (Fig. 2a-c, dashed lines).

**Table 1.**
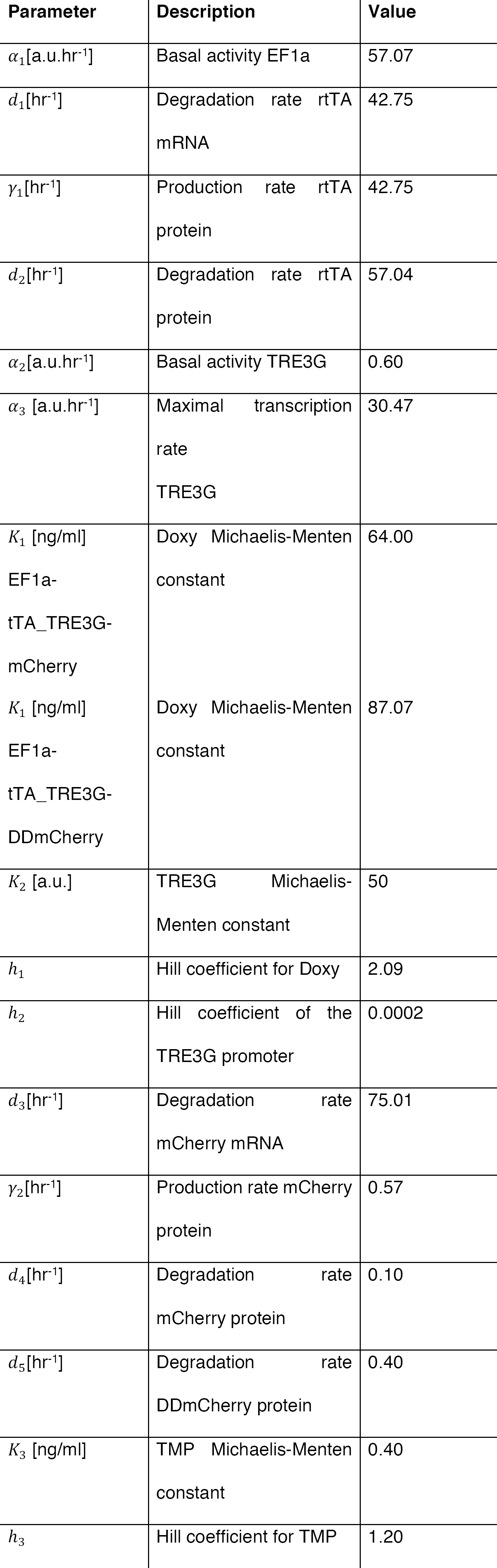
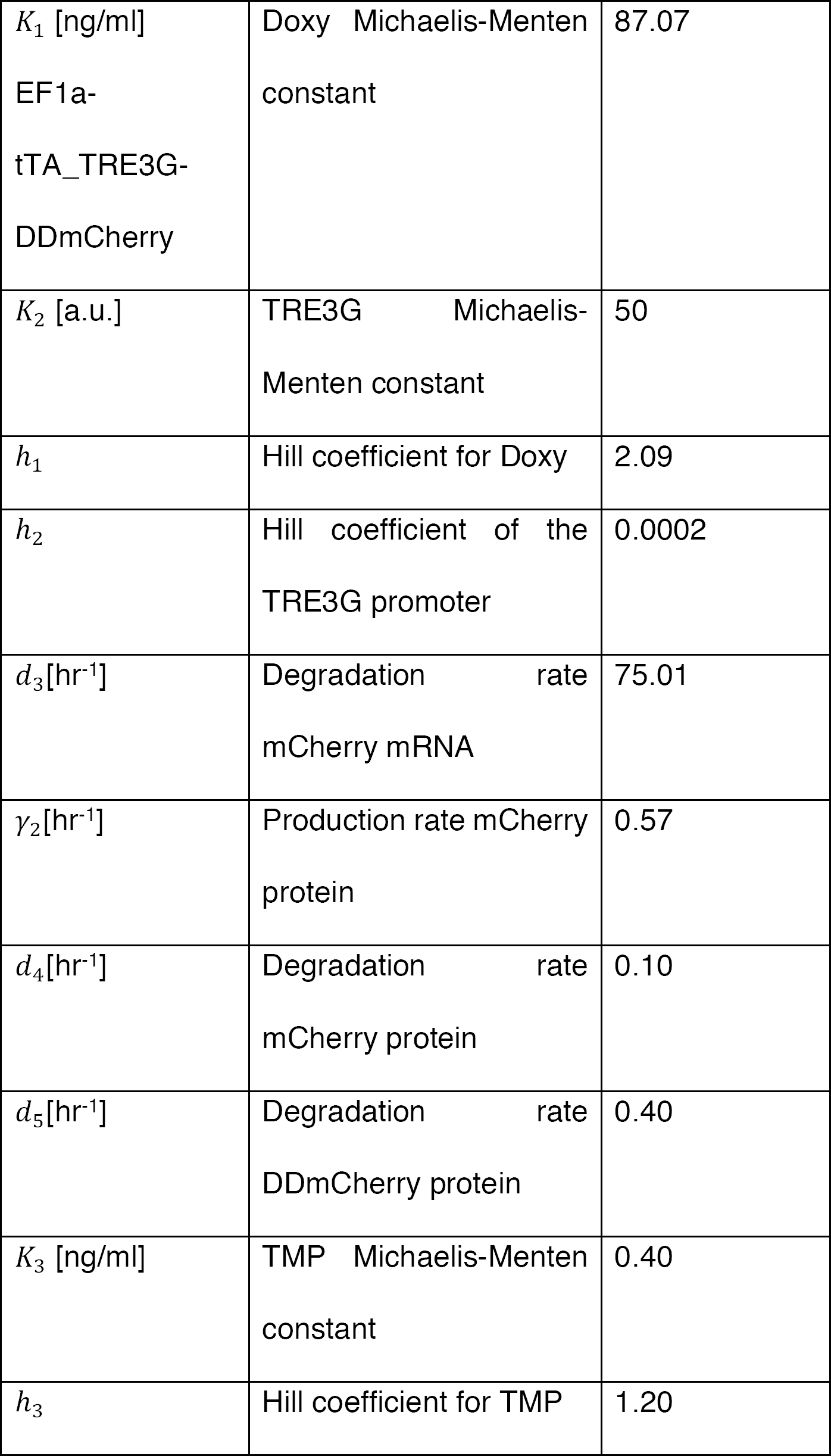
Parameters identified for EF1a-rtTA_TRE3G-mCherry and EF1a-rtTA_TRE3G-DDmCherry systems.

Next, we used the model fitted on titration data to predict the switch-off dynamic response of both the Tet-On and the Tet-On/DD systems upon inducer withdrawal; model simulations indicated much faster switch-off of the dual-input system (Fig. 2d, e, dashed lines). These predictions were validated experimentally: both EF1a-rtTA_TRE3G-mCherry and EF1a-rtTA_TRE3G-DDmCherry mESCs were able to reach full steady-state induction upon 14hrs of incubation with Doxy and Doxy/TMP, respectively, with the conditionally destabilised protein showing 80% protein reduction 8hrs after inducer removal, as compared to 40% reduction only in the Tet on system (Fig. 2d, e, dots; Supplementary Fig. 2c, d).

These results indicate that the dual-input system we developed allows fully tunable protein induction with both drugs, and faster switch-off dynamics as compared to a standard Tet system. Furthermore, the ODE model, fitted on steady-state data, satisfactory replicated experimental time-course dynamics, indicating that Hill kinetics suit modelling Destabilising Domain responses, for which a mathematical formalism has been missing.

### Modular control and dynamic range of transcriptional activation and protein stability

Given the improved dynamic response of the conditionally destabilised mCherry we measured in EF1a-rtTA_TRE3G-DDmCherry mESCs, we reasoned about further exploiting the performance of the inducible system by fusing the rtTA with a Destabilising Domain. Therefore, we generated two additional mESC lines constitutively expressing a conditionally destabilised version of the rtTA in combination with either a stable or a conditionally destabilised mCherry (Supplementary Fig. 2e, f; EF1a-DDrtTA_TRE3G-mCherry and EF1a-DDrtTA_TRE3G-DDmCherry, respectively).

Both cells lines responded, as expected, to Doxy/TMP treatment at steady-state, and Doxy and TMP titration experiments indicated again good response of mCherry to both drugs (Supplementary Fig. 2e-j, m and n). Interestingly, when DD is present on both the transactivator (rtTA) and the fluorescent reporter, the residual protein stability when the mRNA is transcribed is significantly reduced (compare +Doxy/-TMP samples in Supplementary Fig. 2b and Supplementary Fig. 2n).

The fitting of two new sets of ODEs, describing EF1a-DDrtTA_TRE3G-mCherry and EF1a-DDrtTA_TRE3G-DDmCherry systems consistently with the aforementioned modelling assumptions (Supplementary Information), showed good agreement with experimental data (Supplementary Fig. 2g-j, dashed lines and dots represent model fitting and experimental data, respectively).

Performing time-course experiments as for EF1a-rtTA_TRE3G-DDmCherry mESCs, we did not observe any further improvement in protein switch-off kinetics upon inducer removal (compare Fig. 2e and Supplementary Fig. 2k, l, dots; corresponding data of the latter figures in Supplementary Fig. 2o, p). Validation of the models on time-course data (Supplementary Fig. 2k, l, dashed lines) also confirmed these results.

We argue that the observed switch-off dynamics are caused by a partial persistence of protein stability despite TMP removal, as also previously shown by blocking protein synthesis in EF1a-rtTA_TRE3G-DDmCherry mESCs in absence of TMP (Fig. 1g; Supplementary Fig. 1b). Furthermore, EF1a-DDrtTA_TRE3G-DDmCherry mESCs also showed reduced activation in presence of both Doxy and TMP (compare Supplementary Fig. 2b and n), possibly due to limited TMP efficacy in stabilizing both DDrtTA and DDmCherry.

Nevertheless, the dual-input control system allows fully tunable dynamic range of protein levels, as shown by dose-response model simulations (Fig. 2f; Supplementary Fig. 2q, s). We simulated steady-state cellular response to different combinations of both Doxy and TMP in EF1a-rtTA_TRE3G-DDmCherry, EF1a-DDrtTA_TRE3G-mCherry and EF1a-DDrtTA_TRE3G-DDmCherry mESCs and found a robust dose-response increase in mCherry protein levels (Fig. 2f; Supplementary Fig. 2q, s). Experimental validation of model simulations (Fig. 2g; Supplementary Fig. 2r, t) further confirmed both validity of the models, and the ability of tightly controlling the exogenous protein dynamic range when modulating both drugs.

### Dual-input engineering of gene expression dynamics using *in silico* feedback control

Recently, successful attempts have been made to dynamically control and regulate gene expression patterns in living cells, applying control engineering paradigms and using microfluidics or optogenetics platforms for dynamic cell stimulation, and microscopy or flow cytometry for obtaining real-time cell read-outs.^28-37^. Applications in mammalian cells have been recently attempted^29^, controlling a Tet-Off promoter-driven fluorescent protein using tetracycline as control input; this pioneering work showed feasibility of the *in silico* feedback control action, but highlighted challenges in controlling a system with slow kinetic response.

We wondered if our dual-input system would allow finer *in silico* gene expression - feedback control using a microfluidic/microscopy platform (Supplementary Fig. 3); thus, we tested the effectiveness of individual gene transcription or protein stability control regulation as compared to the combined modulation. EF1a-rtTA_TRE3G-DDmCherry mESCs, Doxy/TMP treated, showed the same activation profile of EF1a-rtTA_TRE3G-Cherry mESCs (Supplementary Fig. 2a, b, Doxy and Doxy/TMP respectively); thus, *in silico* feedback control experiments were performed on EF1a-rtTA_TRE3G-DDmCherry mESCs only using different combination of control inputs. Specifically, for set-point control experiments (Supplementary Fig. 3), we applied a Relay control strategy to reduce protein expression to 50% of the maximal induction (set-point, control reference)^29^. Control inputs were: time-varying Doxy (Fig. 3a), TMP (Fig. 3b), and Doxy/TMP (Fig. 3c).

**Figure 3.**
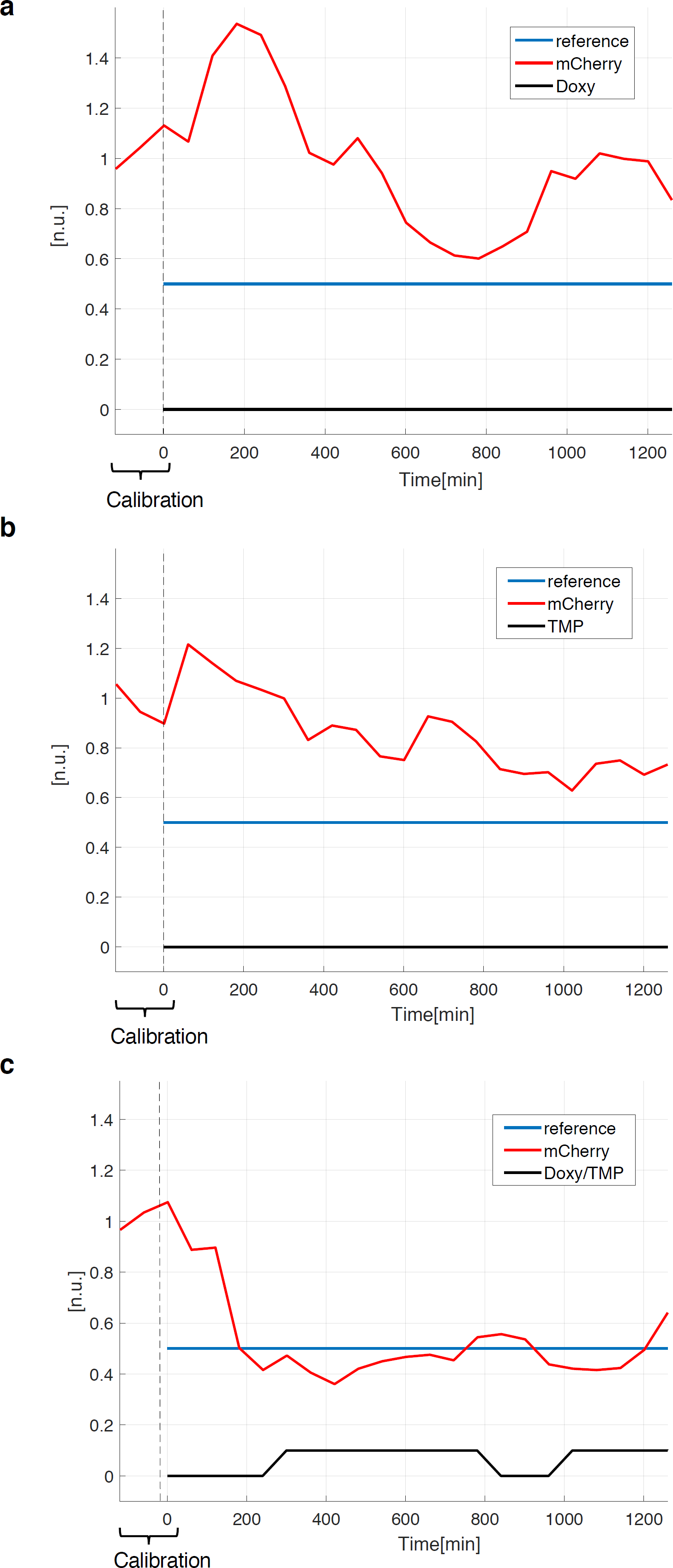
*In silico* feedback control of gene expression in mESCs. **(a-c)** Set-point control experiment, using inducers (Doxy 1000ng/mL and/or TMP 100nM) as control inputs. In red the measured output (normalised mCherry fluorescence), in blue the control reference fluorescence, set at 50% of the average values measured during the calibration phase (120mins with continuous Doxy/TMP administration). The time-lapse sampling time was set at 60mins. The effectiveness of controlling transcription **(a)**, protein stability **(b)** and both **(c)** was tested: cells received either media with both inducers **(a-c)** when the measured fluorescence is below the control reference, and plain media **(c)** or Doxy **(a)** or TMP **(b)** supplemented media when the measured fluorescence is above the control reference. Details about microfluidics device, segmentation and control algorithms, and live cell imaging setting, are provided in Supplementary Information.

We found that the control action fails in controlling DDmCherry levels to reach and maintained the set-point when only gene transcription (Fig. 3a) or protein stability (Fig. 3b) are modulated, whereas a vastly improved control is achieved when both TMP and Doxy are applied (Fig. 3c). Of note, cells are able to reach the set-point with much faster dynamics than the ones previously observed using a Tet-Off system^29^.

These results demonstrate that our dual-input system is preferable for *in silico* feedback control of gene expression in mammalian cells, and is likely to be suitable for generating more complex time varying gene-expression dynamics.

### Dual-input control of signalling pathway in embryonic stem cells

We then tested the general applicability our dual-input system for fine-tune modulation of signalling pathways. We chose the canonical Wnt signalling pathway, as its activation levels and spatiotemporal dynamics are crucial for several biological processes^38,39^, including intestinal crypt homeostasis^40-42^, somite development during embryogenesis^43^, stem cell pluripotency maintenance and somatic cell reprogramming^44,45^.

To test the ability of our system to modulate levels of β-catenin, the main effector of the canonical Wnt pathway^39,46^, we generated mESCs stably expressing a fusion protein comprising the destabilising domain (DD), the mCherry fluorescent protein and a constitutively active β-catenin (β-catenin^S33Y^)^47^, driven by a doxycycline-inducible promoter (henceforth EF1a-rtTA_TRE3G-DDmCherryβ-catenin^S33Y^ mESCs, Fig. 4a). As with previous constructs, we found selective response of exogenous β-catenin^S33Y^ mRNA (Supplementary Fig. 4a, b) and protein (Fig. 4a-d; Supplementary Fig. 4c, d) levels to Doxy and Doxy/TMP treatment, respectively. To prove functionality of the recombinant protein, we measured the subcellular localisation of both endogenous and exogenous forms of β-catenin and found comparable patterns upon inducer treatment (Fig. 4c, d; Supplementary Fig. 4c, d). Furthermore, nuclear translocation of the exogenous protein was detected from western-blot on subcellular fractions (Supplementary Fig. 4e), confirming full functionality of the conditional protein.

**Figure 4.**
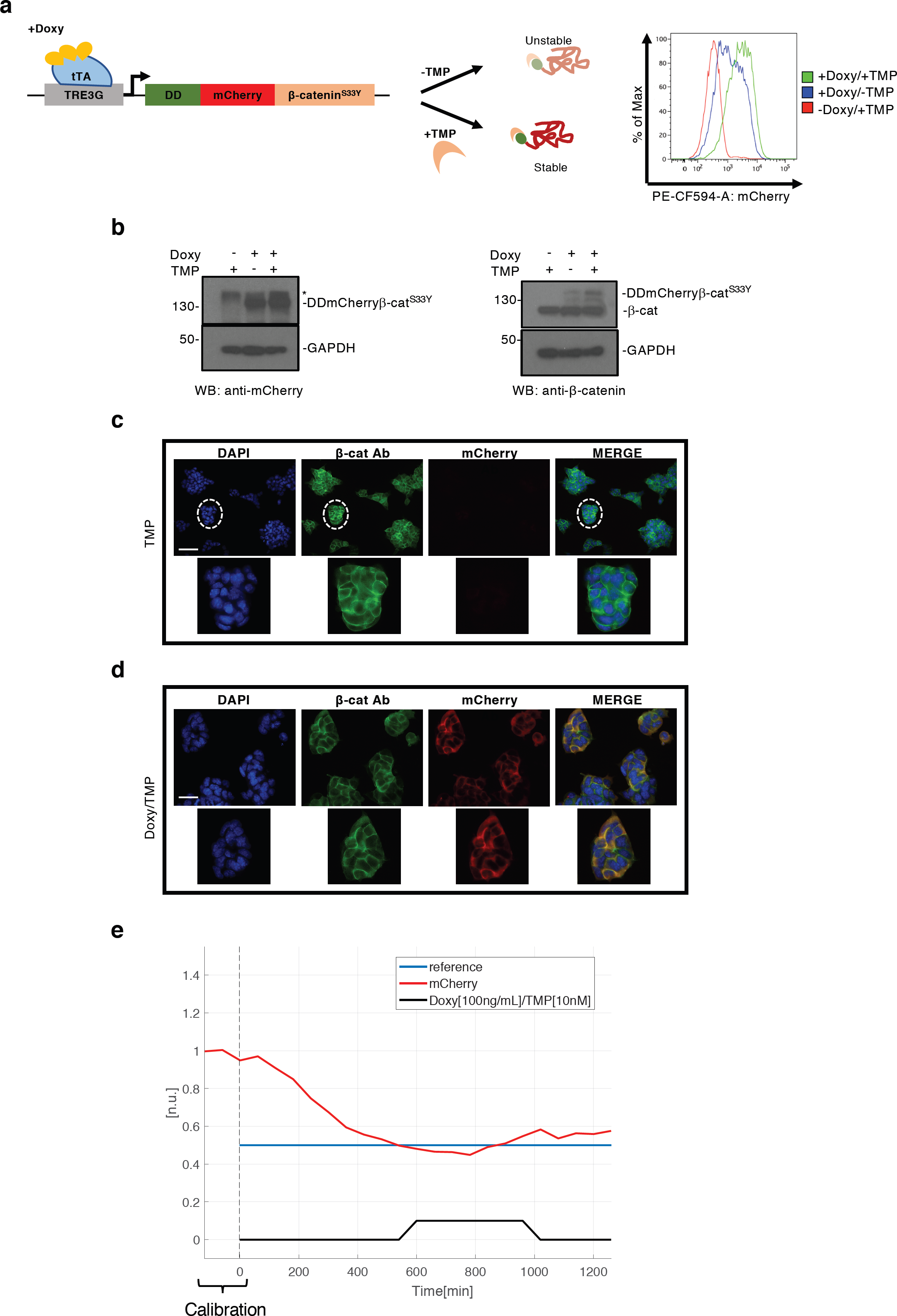
Dual-input control of β-catenin levels in mESCs. (**a, left panel**) Cartoon of the designed dual-input control system consisting of a reverse transactivator (rtTA) and conditionally destabilised and constitutively active β-catenin (β-catenin^S33Y^), fused to the mCherry fluorescent protein. **(a, right panel)** Representative flow cytometry profile of Doxy and/or TMP treated EF1a-rtTA_TRE3G-DDmCherryβ-catenin^S33Y^mESCs. **(b)** DDmCherryβ-catenin^S33Y^protein levels measured by western-blot in EF1a-rtTA_TRE3G-DDmCherryβ-catenin^S33Y^ mESCs treated with Doxy and/or TMP; mCherry (**left panel**) and β-catenin (**right panel**) antibodies were used. * indicates a non-specific band. **(c, d)** β-catenin immunostaining in EF1a-rtTA_TRE3G-DDmCherryβ-catenin^S33Y^ mESCs treated for 24hrs with TMP **(c)** or Doxy/TMP **(d)**, using mCherry (red signal) and β-catenin (green signal) antibodies. DAPI was used to stain the nuclei. Zoomed pictures of selected clones are shown. Scale bars 25μm. **(a-d)** Doxy and TMP were used at 1000ng/mL and 100nM, respectively. **(e)** Set-point control experiment, using inducers (Doxy 100ng/mL and TMP 10nM) as control inputs. In red the measured output (normalised mCherry fluorescence), in blue the reference fluorescence, set at 50% of the average values measured during the calibration phase (120mins with continuous Doxy/TMP administration). The time-lapse sampling time was set at 60mins. Cells received either media with both inducers when the measured fluorescence is below the control reference or plain media when the measured fluorescence is above the control reference. Details about microfluidics device, segmentation and control algorithms, and live cell imaging setting, are provided in Supplementary Information.

We tested DDmCherryβ-catenin^S33Y^ tunability by running *in silico* feedback control experiments, using Doxy/TMP as time-varying inputs as they allowed robust set-point regulation in DDmCherry mESCs (Fig. 3c). In presence of maximal concentrations of Doxy (1000ng/mL) and TMP (100nM), DDmCherryβ-catenin^S33Y^ protein levels were stable and never reached the desired set-point (Supplementary Fig. 4f), in contrast with the control experiment in Fig. 3c, likely due to different protein half-life and size of DDmCherry as compared to DDmCherryβ-catenin^S33Y^. Therefore, we performed titration experiments of EF1a-rtTA_TRE3G-DDmCherryβ-catenin^S33Y^ mESCs (Supplementary Fig. 4g) and consequently selected lower concentrations of Doxy (100ng/mL) and TMP (10nM), still enabling full activation but possibly easier to be washed out from cells, to run set-point control experiments. Lower concentration of inducer molecules allowed robust feedback control (Fig. 4e), demonstrating that control of biologically-relevant proteins is possible, and that, irrespectively of half-life and protein turnover, our system is able to fine tune gene-of-interest levels.

These findings open potential new avenues for dynamic control of protein expression in stem cells that may help addressing key questions of how dynamic patterns of Wnt pathway activation can direct cell fate choices.

## Discussion

Complex time-varying and dynamic patterns of gene expression underlie numerous biological processes such as immune regulation and cell fate choice^48, 49^;mimicking and perturbing these dynamical profiles of expression is technically challenging and requires tools that faithfully recapitulate often fluctuating kinetics.

In this study, we presented an enhanced tool for conditional gene expression encompassing a Tet-on 3G system combined with DD technology that allows fast, specific, reversible and tunable perturbation of biological macromolecules. We demonstrated experimentally and with a computational model that by incorporating control at the transcriptional level, as well as regulating protein-of-interest stability, allows a much finer and tighter control of gene expression kinetics compared to transcriptional (Tet-on) control alone. Indeed, the key advantage of our system is that genes of interest can be robustly induced but also much more rapidly switched off, likely by enhanced proteasomal degradation of the translated protein, compared to either stand-alone transcriptional or post-translational control.

Inducible promoters enable transcriptional regulation and, combined with additional systems for protein levels modulation, provide valuable tools for controlling the expression rate and amount of proteins of interest. The destabilisation domain we applied in this study, although being slower in switch-off dynamics than Auxin-inducible degradation, overcomes the use of hormones, which can vary across different biological systems, and requires minimal genetic manipulations^18,19^. Furthermore, as the inducer stabilises the protein, the dynamic range of the inducible system can be fully controlled to obtain ‘on demand’ activation levels.

We demonstrated the applicability of our dual-input regulation system for microfluidics/microscopy-based *in silico* feedback control, showing that it outperforms transcriptional- and post-translational-only manipulation to reach and maintain a set-point control reference, even using a simple and model-free control strategy.

Our system suits generation of both spatial- and temporal-patterns of gene expression that traditionally have not been easy to manipulate. For instance, the Wnt/β-catenin pathway is known to display oscillating patterns of gene expression in stem cells and in the developing embryo^43-45^, which are a key determinant of cell fate determination. We demonstrated that finely tuned levels of β-catenin are achievable, despite the half-life of the tagged protein. This suggests that more complex dynamic patterns of Wnt-pathway activation could be engineered; mimicking *in silico* native temporal patterns should be compatible with the ON-OFF kinetics we measured, given the relatively slow period and amplitude of oscillations observed *in vivo*^*43*^. Our platform, which has the potential to allow a quantitative assessment of the molecular mechanisms and dynamics underpinning cell fate choices, paves the way for controlling cellular behaviour in clinically-relevant model systems, such as stem cells.

The inducibility and switch-like behaviour of our system makes it a powerful perturbation method, and the ability to tune protein levels with a high degree of predictability (as evidenced by our faithful capture of system dynamics using a computational model) makes it a valuable resource with a broad scope of applications. For instance, it could be harnessed to study how signalling networks or synthetic gene circuits are wired together, by fine manipulation of one ‘node’ in the network and observing how this affects other cognate members within the circuit. Ultimately, this approach could be used to deconvolute and quantitatively assess the interactions within these networks, which could both explain and predict the consequences of a given perturbation. The mathematical model we fitted and validated could be used to test the design of novel synthetic networks based on inducible promoters and destabilising domains^27,50,51^.

Finally, the ability of both Doxy and TMP to cross the placental barrier^52^ opens up the possibility that our dual-input system could allow fine-tuning of genes and signalling pathways essential in embryonic development, and could be applicable as a novel approach for targeted and modulable gene therapy.

## Materials and methods

### Inducible constructs

pLVX_EF1a-Tet3G Neo (in the manuscript referred as EF1a-rtTA) and pLVX_TRE3G Puro were purchased from Clontech (631363). DHFR-derived destabilization domain (DD) and mCherry were PCR amplified from pOddKS^53^ and 7TGC^54^ plasmids, respectively. Following manufacturer’s instructions, fragments were assembled into pLVX_TRE3G Puro (Clontech) and linearized with SmaI (NEB) using HiFi-DNA assembly cloning kit (NEB). pLVX_TRE3G-DDmCherry is available on Addgene (Plasmid #108679).

### Cell line derivation

Stable cell lines were generated by lentiviral infection of R1 (EF1a-rtTA_TRE3G-mCherry, EF1a-rtTA_TRE3G-DDmCherry TET, EF1a-DDrtTA_TRE3G-mCherry and EF1a-DDrtTA_TRE3G-DDmCherry) and E14Tg2a (EF1a-rtTA_TRE3G-DDmCherryβ-catenin^S33Y^) mouse embryonic stem cells (mESCs) as in^55^. mESCs were firstly infected with the stable (EF1a-rtTA) or conditionally destabilised (EF1a-DDrtTA) transactivator containing vector. After Neomycin selection, cells were subjected to a second round of infection with the doxycycline-inducible vector (pLVX_TRE3GmCherry, DDmCherry or DDmCherryBcatS33Y) and Puromycin selected. Finally, to enrich for mCherry expressing cells and to homogenise mCherry levels across cell lines, we sorted mCherry positive cells after Doxy or Doxy/TMP (1000ng/mL and 10μM, respectively) 24hrs treatment. mESCs were cultured on gelatin coated dishes in knockout Dulbecco’s modified Eagle’s medium (DMEM) supplemented with 20% fetal bovine serum (Sigma), 1 x nonessential amino acids, 1 x GlutaMax, 1 x 2-mercaptoethanol, and 1000 U/ml LIF (Peprotech).

### Flow activated cell sorting (FACS)

Cells were washed with sterile Phosphate-Buffered Saline (PBS, Gibco), trypsinised for 2-3’ at room temperature and centrifuged at 1000rmp for 5’. Pelleted cells were resuspended in 500uL of complete mESCs media supplemented with DAPI. The mCherry positive fraction was sorted from DAPI negative using the BD Influx high-speed 16-parameter fluorescence activated cell sorter; the gating strategy was defined to get comparable mCherry intensity across cell lines.

### Flow cytometry analysis

Cells from a 24well plate were washed with sterile Phosphate-Buffered Saline (PBS, Gibco), incubated with 50μL of trypsin for 2-3’ at room temperature and collected with 150μL of PBS 2% FBS containing DAPI as cell viability marker. Cell suspension was analysed using the BD LSR Fortessa and 10000 living cells were recorded for each sample. Both % of mCherry positive cells and Median Fluorescence Intensity (MFI) were calculated over living cells, gated as DAPI negative.

### Drug treatments

Drugs used in this study are doxycycline (Sigma, D9891-1G), TMP (Sigma, T7883), cycloheximide (Sigma, C4859) and MG132 (Tocris, 1748). Concentrations and time of treatment are specified in main text and figure legends.

### qPCR

For quantitative PCR, the total RNA was extracted from cells using the RNeasy kit (Qiagen), and the cDNA was generated from 1μg of RNA. Twenty-five ng of cDNA were used as template for each qPCR reaction, in a 15μl reaction volume. iTaq Universal SYBR Green Supermix (1725120, Bio-Rad) was used with Qiagen Rotor-Gene System.

### Primers

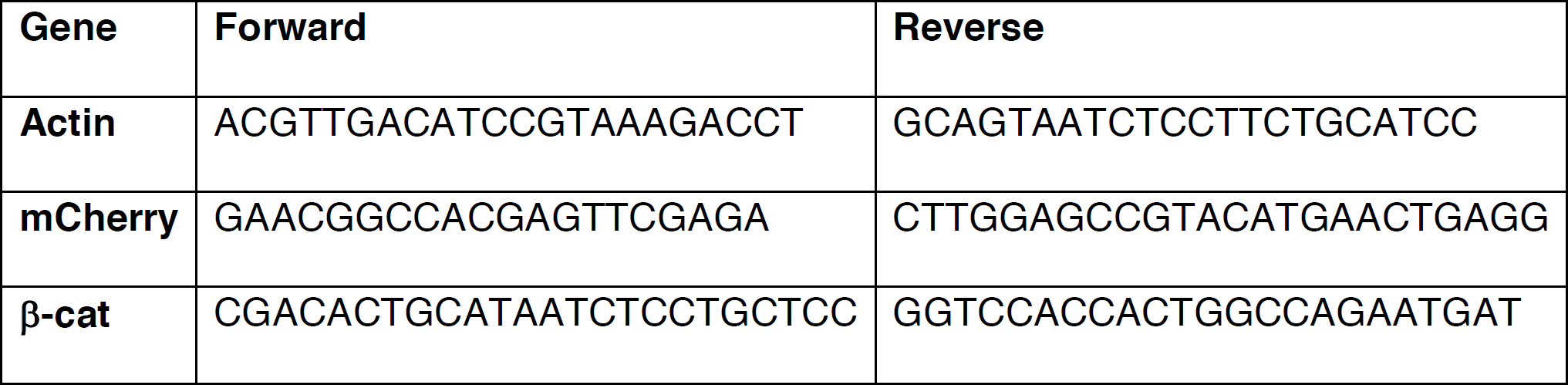

### Biochemical analysis

mESC cell lines were treated as described in the text and figure legends; Doxy and TMP are specified in the latter; cycloheximide was used at 25μg/ml. Cells were lysed in lysis buffer (1% TX-100, 150mM NaCl, 15mM Tris pH 7.5 and protease inhibitors) before processing by SDS-PAGE and western-blotting. For proteasomal inhibition experiments, MG132 (5μM) was added 6hrs prior to lysis. Antibodies used are anti-mCherry (AB0040-200, Sicgen, 1:10,000), anti-β-catenin, clone 14 (610153, BD Biosciences, 1:5000) and anti-GAPDH, clone 6C5 (ab8245, Abcam, 1:10,000). Image densitometry was carried out using ImageJ.

### Ubiquitination analysis

Cells were drug treated as described in the text and figure legends, with Doxy (1000ng/mL) and TMP (100nM) for 24hrs, and chased for 12hrs with or without TMP in the presence of MG132. Cells were then lysed in denaturing lysis buffer (2% SDS, 0.5 % NP40, 15mM Tris pH 6.8, 5% glycerol, 150mM NaCl) to prevent deubiquitination, before dilution in modified RIPA buffer (25mM Tris pH7.2, 150mM NaCl, 1% NP40, 0.5 % Sodium Deoxycholate, 1mM EDTA) to dilute the SDS to 0.1%. Samples were then immunoprecipitated with 2μg of anti-ubiquitn antibody (P4D1, Cell Signaling) for 4hrs before addition of Protein G for a further 1hr, followed by extensive washing and processing via SDS-PAGE and western-blotting with anti-mCherry antibody.

### Immunostaining

Cells were washed twice with phosphate-buffered saline (PBS), fixed with 4% paraformaldehyde for 15mins at RT, and blocked in PBS with 0.1% Triton X-100 and 1% BSA, for 60mins. Cells were incubated overnight with the specific primary antibodies and PBS washed (5’ each wash) before being incubated with the secondary antibody conjugated with fluorescein for 1h at room temperature. Finally, cells were washed and mounted on slides with a few drops of Vectashield, with DAPI (Vector Laboratories). The primary antibodies used for immunostaining are: anti-β-catenin, clone 14 (610153, BD Biosciences), working concentration 1:1000; anti-mCherry (AB0040-200, Sicgen) working concentration 1:1000. The fluorescent conjugated secondary antibodies were diluted 1:1000.

### Nucleus/Cytosol fractionation

For Nuclear and Cytosolic protein fractionation, cells were processed as in^56^.

### Statistical Analysis

Differences between groups were analysed by Student t test. A p-value lower than 0.05 was considered statistically significant.

## Acknowledgements

We thank Prof Austin Smith for the wild-type mouse Embryonic Stem Cells used to generate EF1a-rtTA_TRE3G-DDmCherryβ-catenin^S33Y^ mESCs; Dr Andre Hermann and Dr Lorena Sueiro Ballesteros (Flow Cytometry Facility, University of Bristol), and Dr Mark Jepson and Alan Leard (Wolfson Imaging Facility, University of Bristol) for their support. This work was funded by Medical Research Council grant MR/N021444/1 to LM, by the Engineering and 188 Physical Sciences Research Council grant EP/R041695/1 to LM, and by BrisSynBio, a 189 BBSRC/EPSRC Synthetic Biology Research Centre (BB/L01386X/1) to LM.

## Author Contribution

EP generated inducible cell lines; EP and DLR designed and performed experiments; LP implemented the control strategy; LP, SMO and LM developed the mathematical model; FA generated inducible plasmids; DdB supported microfluidic-based experiments; MPC supported plasmid generation; EP, DLR, LP and LM analysed data; EP, DLR, LP and LM wrote the manuscript; LM supervised the entire project.

---

**Supplementary Figure 1:**
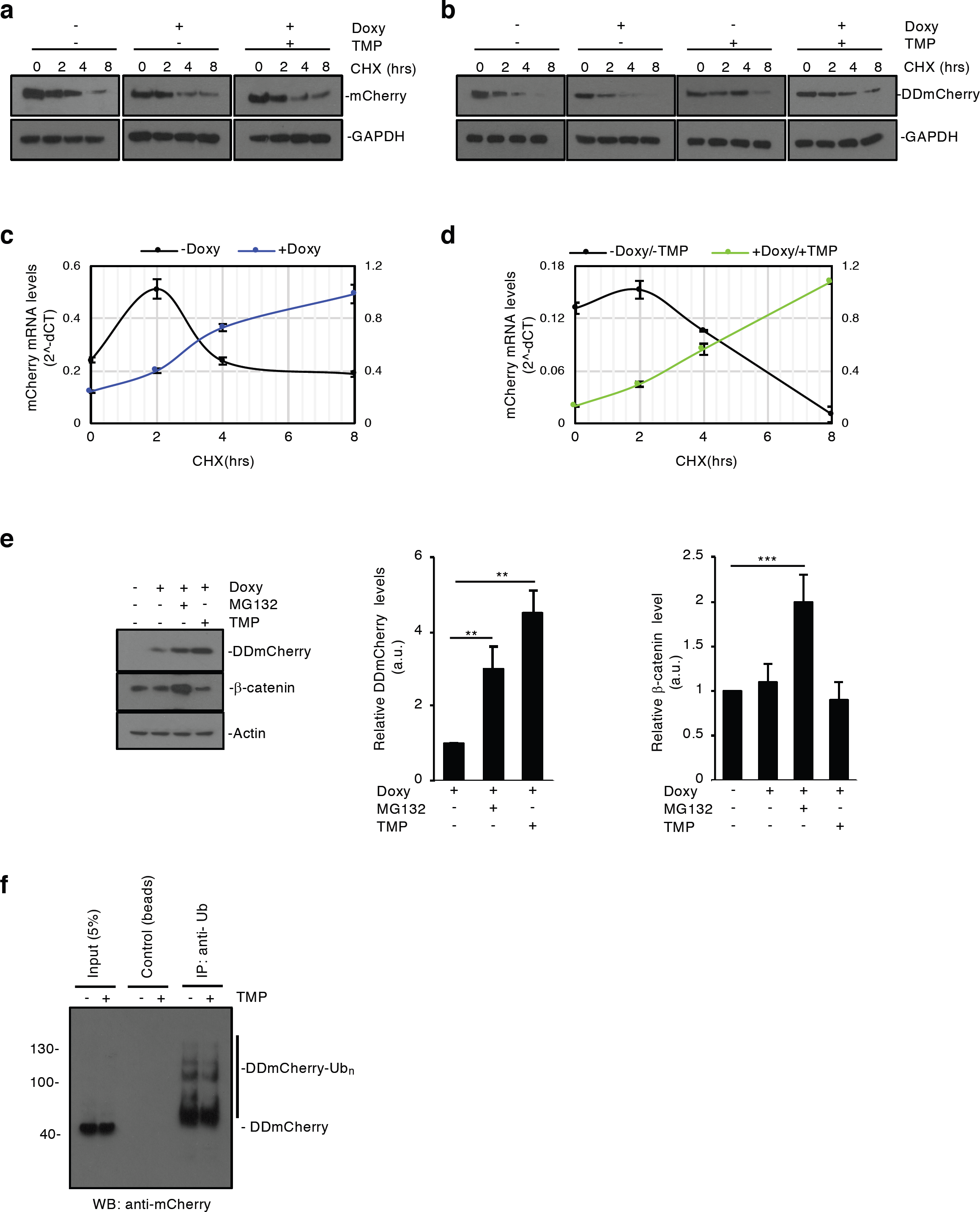
TMP mechanism of action. **(a-d)** mCherry and DDmCherry protein **(a, b)** and mRNA **(c, d)** levels measured by western-blot and qPCR, respectively, in EF1a-rtTA_TRE3G-mCherry **(a, c)** and EF1a-rtTA_TRE3G-DDmCherry **(b, d)**. mESCs were treated as indicated in Fig. 1e and relative legend. **(e)** EF1a-rtTA_TRE3G-DDmCherry mESCs were incubated with Doxy (1000ng/mL) and either TMP (10μM) or MG132 (5μM) for 16hrs. Blocking proteosomal degradation enabled to effectively mimic the stabilising effect of TMP specifically on DDmCherry but not on endogenous β-catenin. Western-blot densitometric quantifications are shown. **(f)** Ubiquitination status analysis. EF1a-rtTA_TRE3G-DDmCherry mESCs were incubated for 24hrs with Doxy (1000ng/mL) and TMP (100nM), washed and incubated with or without TMP (100nM) for additional 12hrs. Samples were processed as indicated in the Materials and methods section, immunoprecipitated with an anti-ubiquitin antibody and blotted with the mCherry antibody to analyse the ubiquitination status of the DDmCherry protein. Note the decrease in poly-ubiquitinated species of DDmCherry when TMP is present. Data are means ± SEM (n=3). p > 0.1, *p < 0.05, **p < 0.01, ***p < 0.0001.

**Supplementary Figure 2.**
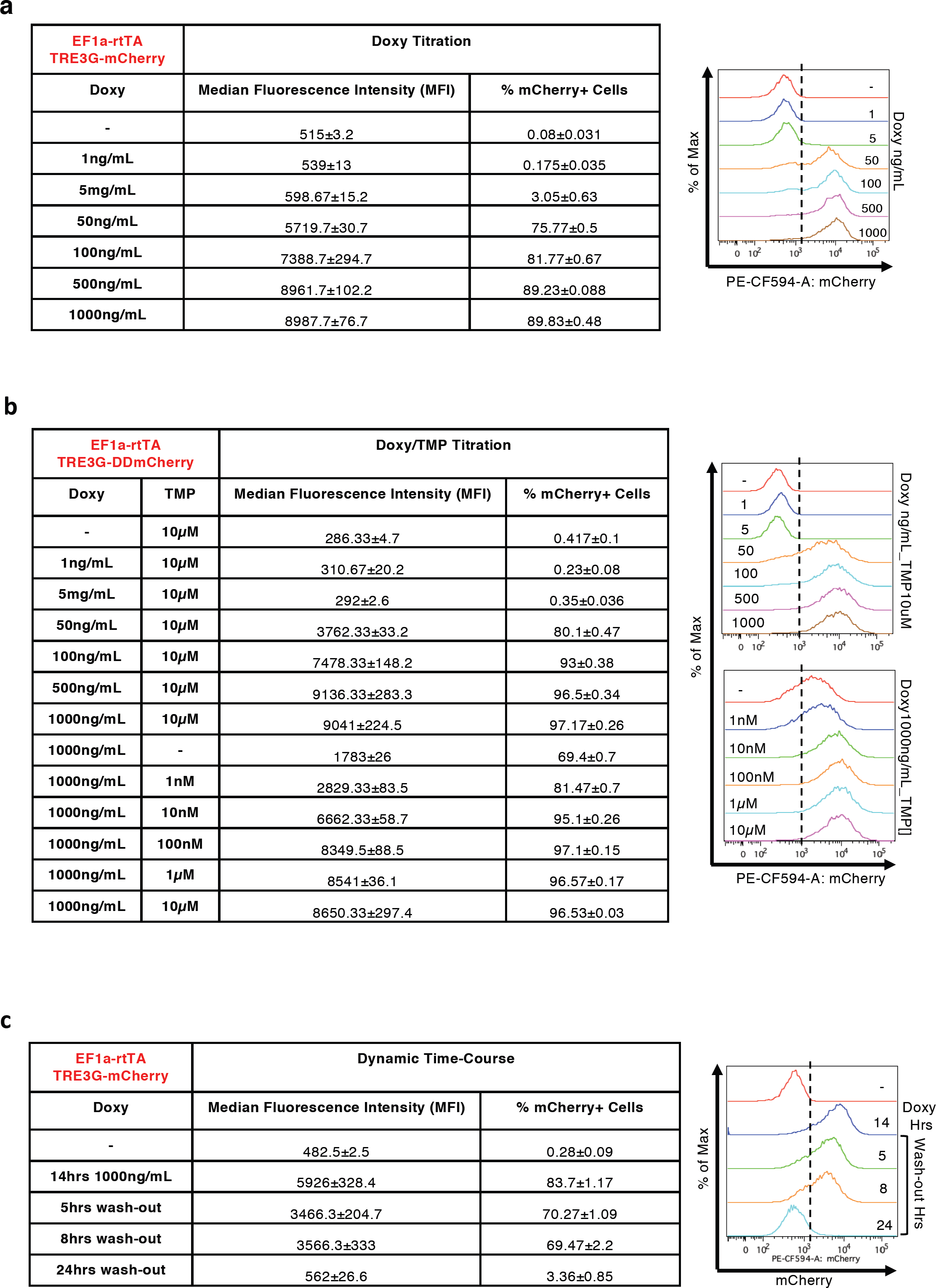
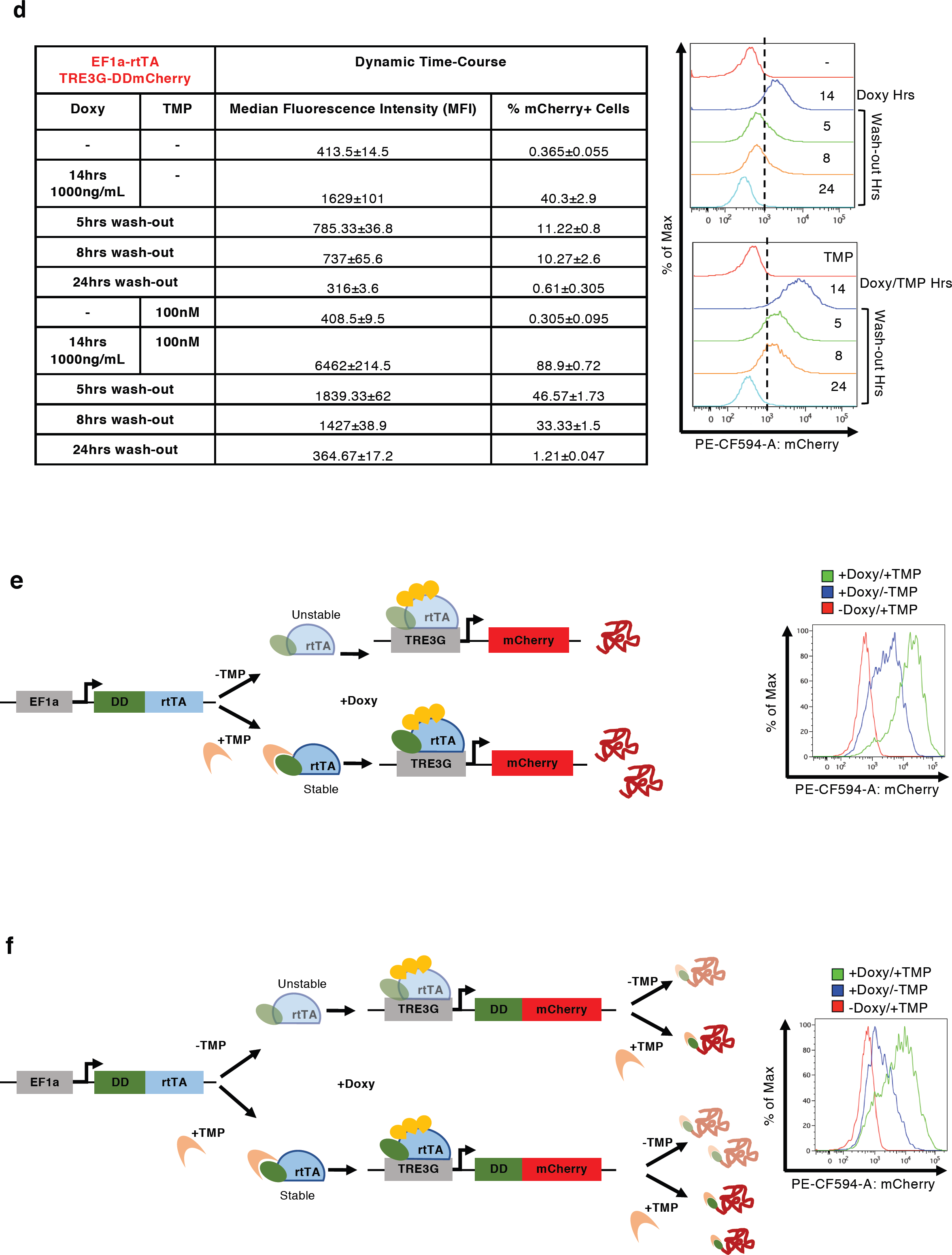
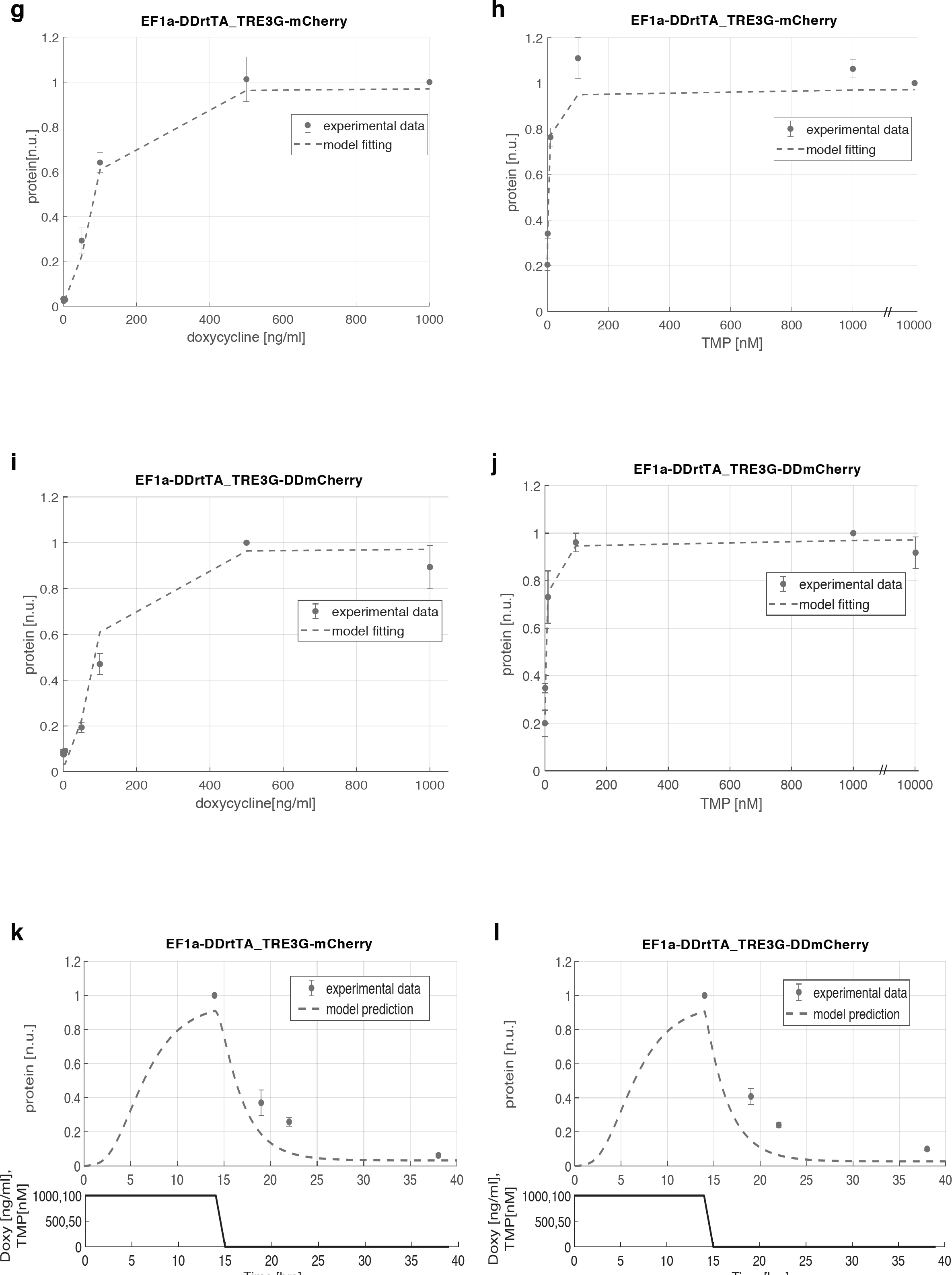
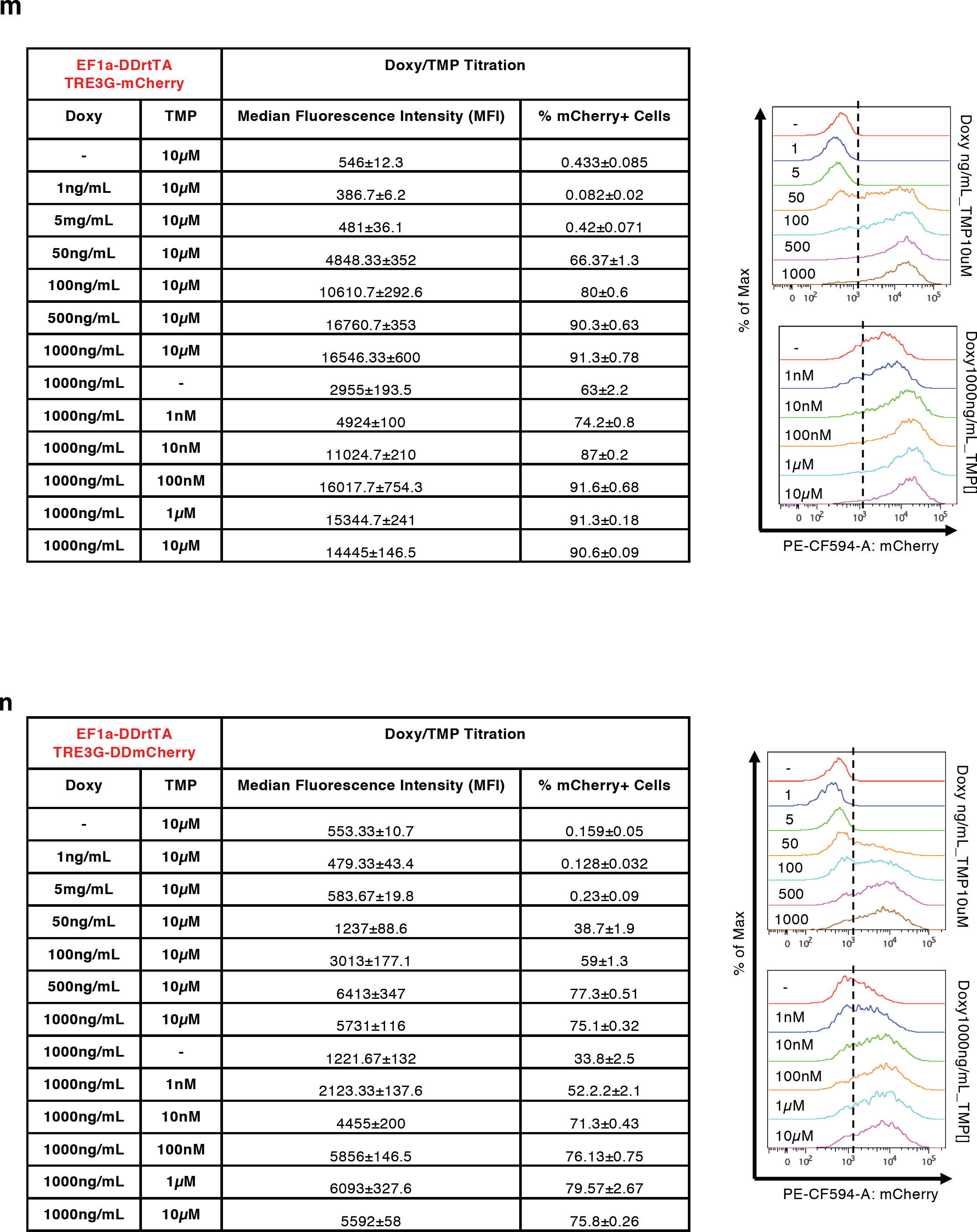
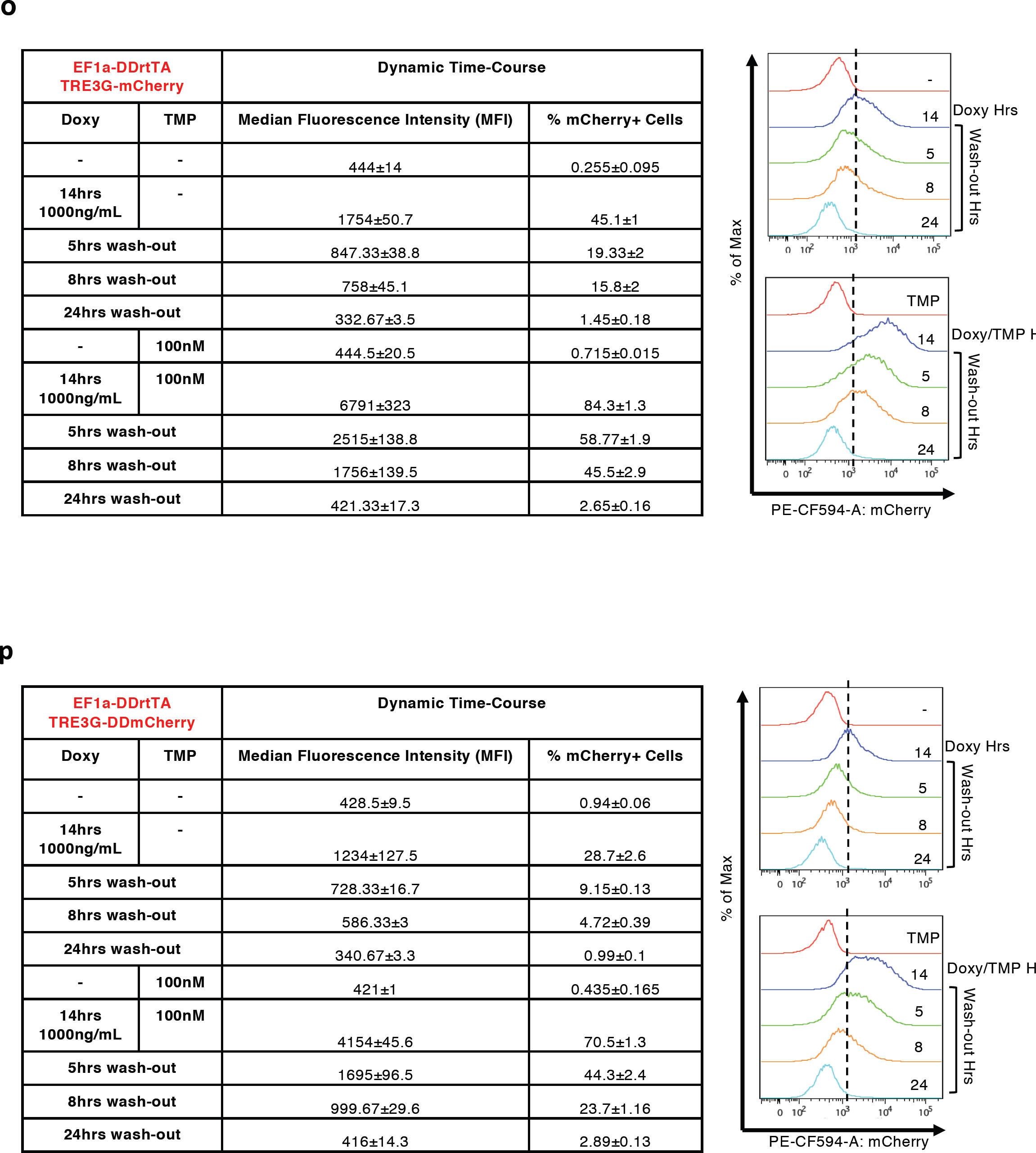
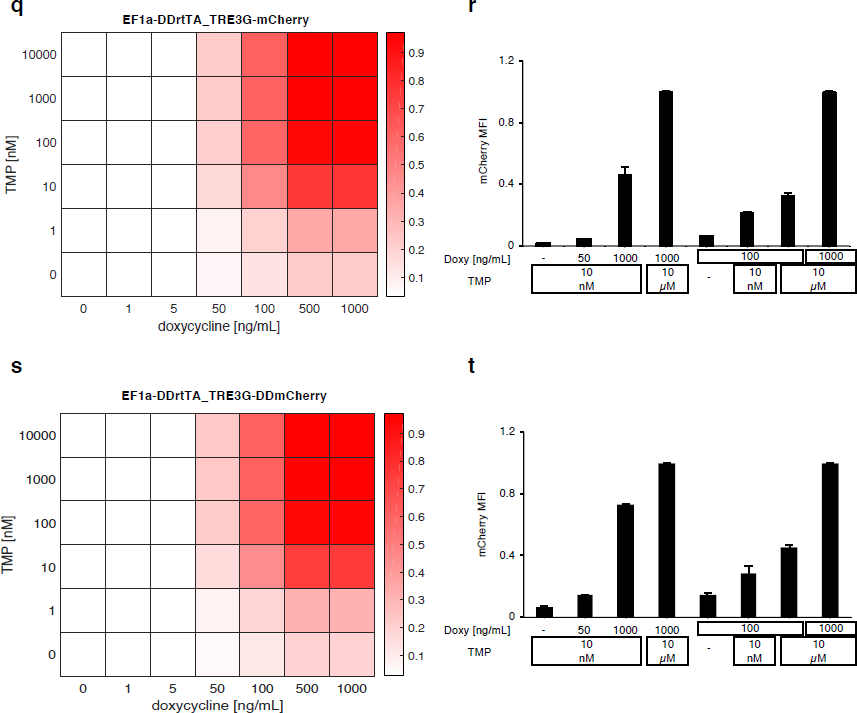
Flow cytometry profiling of inducible mESCs, dose response, switch-off dynamics and dynamic range. **(a-d, m-p)** Median Fluorescence Intensity (MFI) and % of mCherry+ cells measured by flow cytometry in EF1a-rtTA_TRE3G-mCherry **(a, c)**, EF1a-rtTA_TRE3G-DDmCherry **(b, d)**, EF1a-DDrtTA_TRE3G-mCherry **(m, o)** and EF1a-DDrtTA_TRE3G-DDmCherry **(n, p)** mESCs. Inducer titration and dynamic response were performed using concentrations and incubation times indicated in the tables. **(e,f)** Cartoon of the designed dual input system consisting of conditionally destabilised transactivator (DDrtTA) driving a stable **(e)** or a conditionally destabilised **(f)** mCherry fluorescent protein. Protein expression following 24hrs Doxy/TMP treatment (1000ng/mL and 100nM, respectively) in EF1a-DDrtTA_TRE3G-mCherry (**e, right panel)** and EF1a-DDrtTA_TRE3G-DDmCherry **(f, right panel)** mESCs. **(g-j)** Fitted model simulations (dashed lines) and experimental data (dots) of EF1a-DDrtTA_TRE3G-mCherry **(g, h)** and EF1a-DDrtTA_TRE3G-DDmCherry **(i, j)** mESC steady-state response. Dots represent experimental data of MFI in Supplementary Fig. 2m, n, normalised over the maximum activation point. **(k, l)** Model predicted dynamic response (dashed lines) and experimental data (dots) of EF1a-DDrtTA_TRE3G-mCherry **(k)** and EF1a-DDrtTA_TRE3G-DDmCherry **(l)** mESCs. Dots represent experimental data of MFI in Supplementary Fig. 2o, p, normalised over the maximum activation point. **(q-t)** Simulations of the model **(q, s)** and experimental validation **(r, t)** of EF1a-DDrtTA_TRE3G-mCherry **(q, r)** and EF1a-DDrtTA_TRE3G-DDmCherry **(s, t)** mESC steady-state response following 24hrs induction with constant concentration of Doxy (100ng/mL) and varying TMP (0; 10nM; 10μM) or constant TMP (10nM) and varying Doxy (0; 50; 1000ng/mL). Maximum concentrations of Doxy and TMP (1000mg/mL and 10μM, respectively) are used as control. mCherry values are shown as heatmaps of scaled values across the entire dynamical range of expression levels. Data are means ±SEM (n=3, a-d, m-p, r, t); ±SD (n=3, g-l, q, s).

**Supplementary Figure 3.**
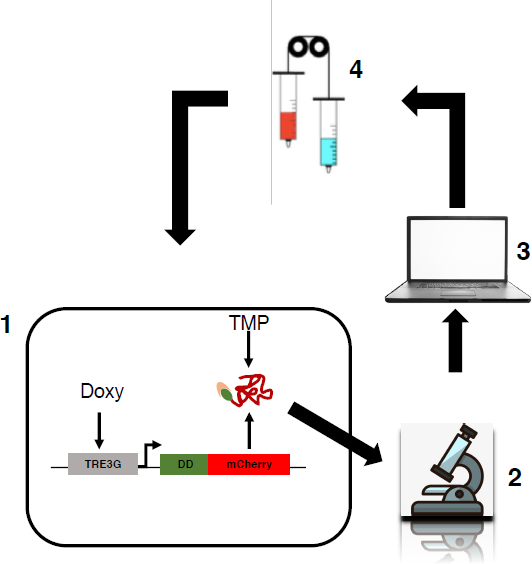
*In silico* feedback control microfluidics/microscopy platform. Schematic representation of the *in silico* feedback control platform used in the present study. It consists of the controlled biological system (1), in our case mouse Embryonic Stem Cells (mESCs) stably carrying an inducible exogenous gene; an inverted fluorescence microscope (2) measuring fluorescence of mCherry and GFP every 60mins during the time-lapse; a computer (3) to implement real-time cell segmentation, fluorescence quantification and a Relay control algorithm (the latter computing the error from the comparison of the desired over the measured and segmented fluorescence); an actuation system (4), consisting of motor-controlled syringes dispensing media +/- inducers (+ inducers if the measured fluorescence is below the set-point and vice versa)^1^.

**Supplementary Figure 4.**
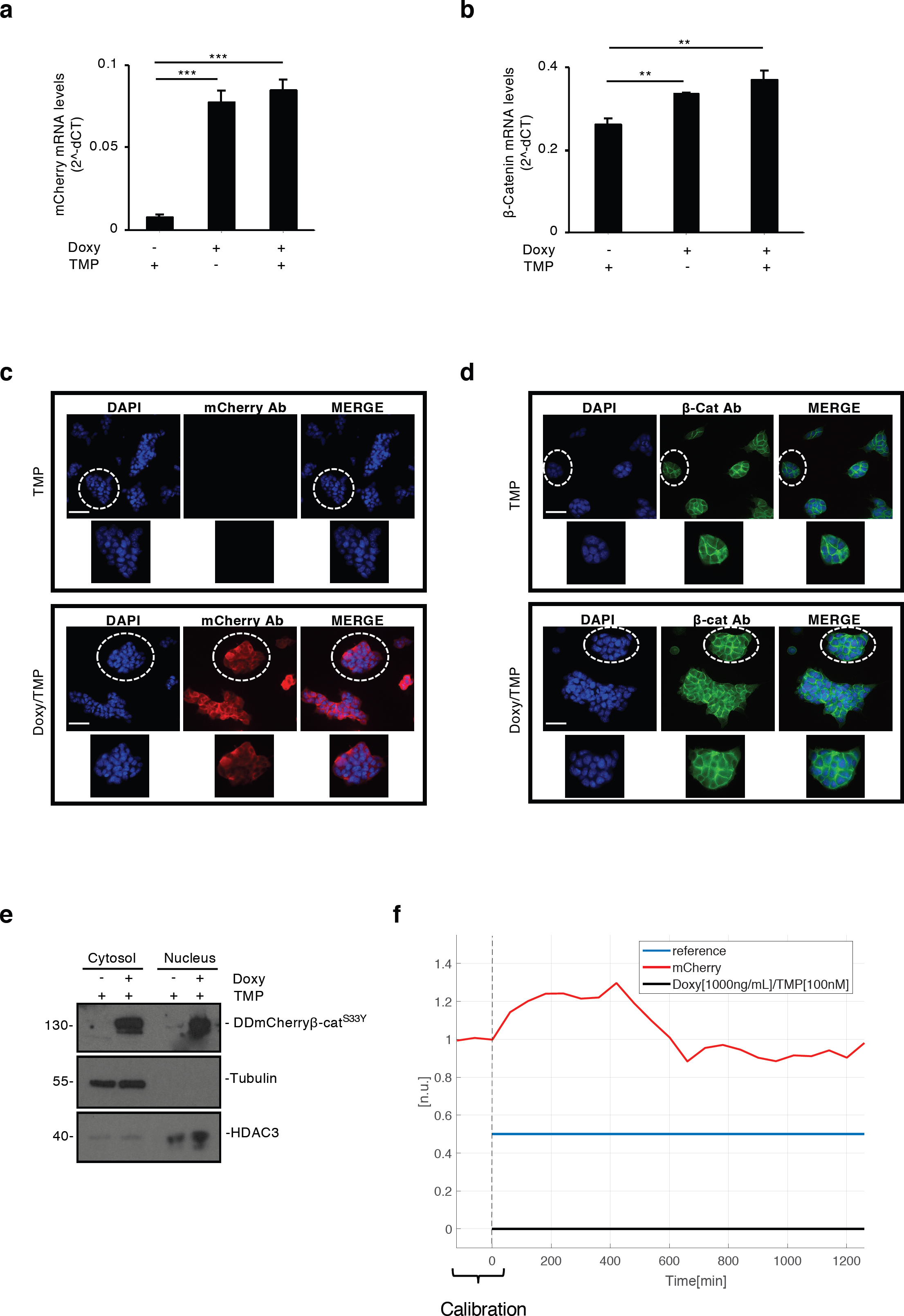
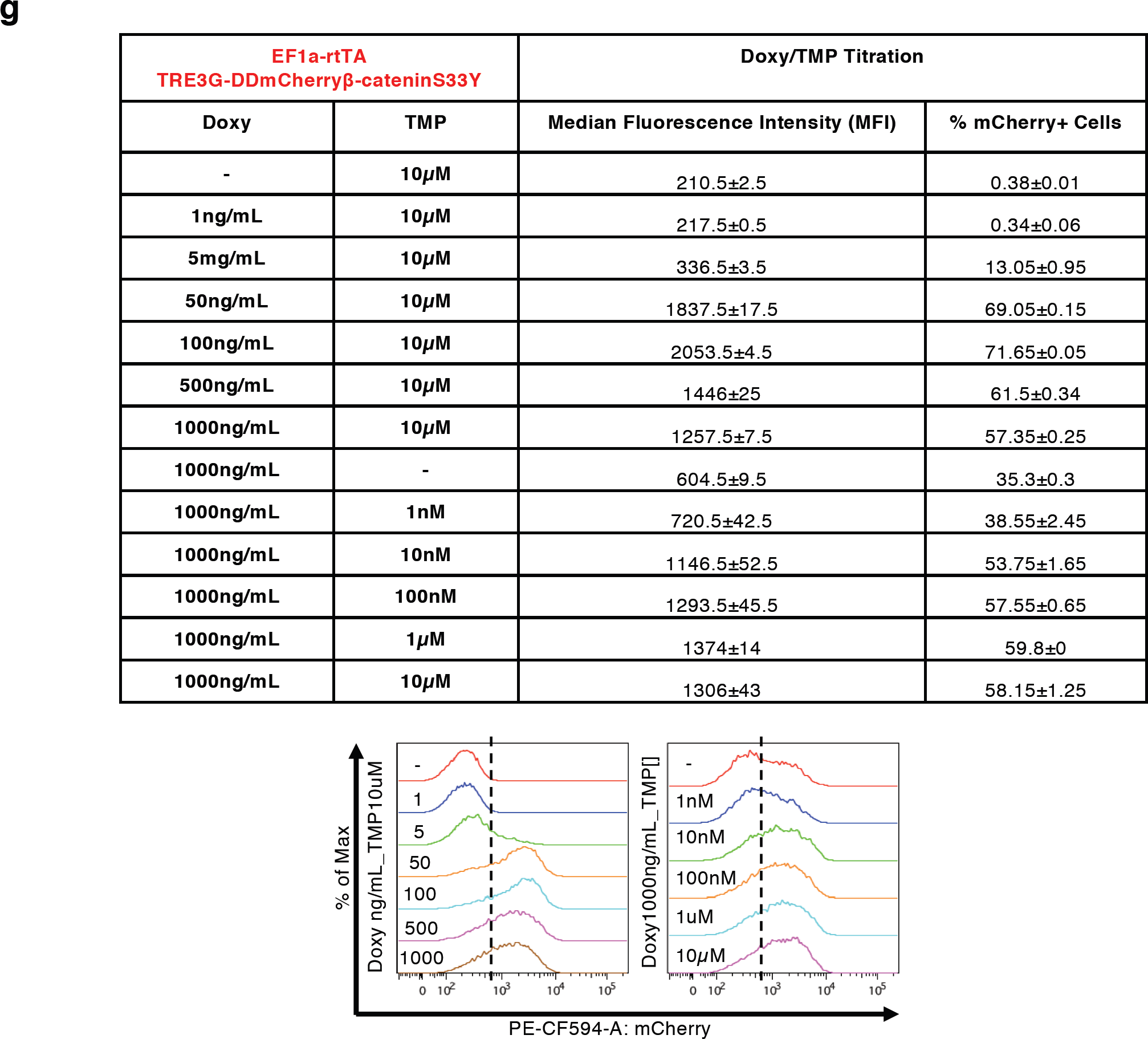
Cell line characterisation and automatic control of β-catenin levels. **(a, b)** Exogenous β-catenin mRNA levels measured by qPCR in EF1a-rtTA_TRE3G-DDmCherryβ-catenin^S33Y^ mESCs induced with Doxy (1000ng/mL) and/or TMP (100nM) for 24hrs. mCherry **(a)** and β-catenin **(b)** specific primers were used. **(c, d)** β-catenin immunostaining in EF1a-rtTA_TRE3G-DDmCherryβ-catenin^S33Y^ mESCs treated for 24hrs with Doxy (1000ng/mL) and/or TMP (100nM); mCherry **(c)** and β-catenin **(d)** antibodies were used. DAPI was used to stain the nuclei. Zoomed pictures of selected cells are shown. Scale bars 25μm. **(e)** Western-blot of nuclear and cytosolic fractions from Doxy/TMP (1000ng/mL and 100nM, respectively) treated EF1a-rtTA_TRE3G-DDmCherryβ-catenin^S33Y^ mESCs, blotted with an anti-mCherry antiboby. **(f)** Set-point control experiment using inducers (Doxy 1000ng/mL and TMP 100nM) as control inputs. In red the measured output (normalised mCherry fluorescence), in blue the control reference fluorescence, set at 50% of the average value measured during the calibration phase (120mins with continuous Doxy/TMP administration). Time-lapse sampling time of 60mins. Cells received either media with both inducers when the measured fluorescence is below the reference, or plain media when the measured fluorescence is above the reference. **(g)** Median Fluorescence Intensity (MFI) and % of mCherry+ cells measured by flow cytometry in EF1a-rtTA_TRE3G-DDmCherryβ-catenin^S33Y^ mESCs after 24hrs of treatment with indicated concentration of Doxy and TMP. Data are means ± SEM (n=2). p > 0.1, *p < 0.05, **p < 0.01, ***p < 0.0001.

## Mathematical modelling

The mathematical models for the EF1a-rtTA_TRE3G-mCherry, EF1a-rtTA_TRE3G-DDmCherry, EF1a-DDrtTA_TRE3G-mCherry and EF1a-DDrtTA_TRE3G-DDmCherry systems are based on sets of 4 ODEs, describing transactivator and fluorescent gene mRNA and correspondent protein concentrations, as the result of a production term and a degradation term. We formulated the parameters estimation problem as a constrained optimization problem:

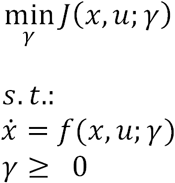

where *x* (is the state vector, *u* is the vector of inputs acting on the systems (e.g. Doxy and TMP), *f* is the vector field of the system dynamics and, *γ* is the vector of all parameter to be identified. *j* is the cost function to be minimized, defined as

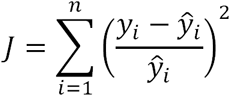

where *n* is the number of experimental data points, *y* are the predicted values of the mathematical model and: *ŷ* are the experimental data points used for fitting. For the parameters identification, we used the Interior Point algorithm implemented in *fmicon* function (Matlab Optimization toolbox ™, Mathworks Matlab R2018a).

The parameters were separately identified for EF1a-rtTA (mCherry and DDmCherry) and EF1a-DDrtTA (mCherry and DDmCherry) mESCs; only the Doxy Michaelis-Menten constant (*K*_1_) was directly fixed from dose-response experimental data. Letting (*x*_1_ be the rtTA mRNA concentration, (*x*_2_ the rtTA protein concentration, (*x*_3_ the mCherry mRNA concentration and (*x* _4_ the mCherry protein concentration, the models of the systems considered in this work are reported below. The description of the estimated parameters and their values are reported in Tables 1 and 2.

**Table 2.** Parameters identified for EF1a-DD::rtTA_TRE3G-mCherry and EF1a-DD::rtTA_TRE3G-DD::mCherry systems.

| Parameter | Description | Value |
| --- | --- | --- |
| $\alpha_1[\text{a.u.hr}^{-1}]$ | Basal activity EF1a | 39.51 |
| $d_1[\text{hr}^{-1}]$ | Degradation rate rtTA mRNA | 91.10 |
| $\gamma_1[\text{hr}^{-1}]$ | Production rate rtTA protein | 39.81 |
| $d_2[\text{hr}^{-1}]$ | Degradation rate rtTA protein | 0.45 |
| $\alpha_2[\text{a.u.hr}^{-1}]$ | Basal activity TRE3G | 0.03 |
| $\alpha_3 [\text{a.u.hr}^{-1}]$ | Maximal transcription rate TRE3G promoter | 2.26 |
| $K_1 [\text{ng/ml}]$<br>EF1a-<br>tTA_TRE3G-mCherry | Doxy Michaelis-Menten constant | 97.45 |
| $K_2 [\text{a.u.}]$ | TRE3G Michaelis-Menten constant | 47.11 |
| $h_1$ | Hill coefficient for Doxy | 2.63 |
| $h_2$ | Hill coefficient of the TRE3G promoter | 3.59 |
| $d_3[\text{hr}^{-1}]$ | Degradation rate mCherry mRNA | 0.41 |
| $\gamma_2 [\text{hr}^{-1}]$ | Production rate mCherry protein | 0.83 |
| $d_4[\text{hr}^{-1}]$ | Degradation rate<br>mCherry protein | 0.1.61 |
| $d_5[\text{hr}^{-1}]$ | Degradation rate<br>DDmCherry protein and<br>DDrtTA protein | 0.31 |
| $K_3 [\text{ng/ml}]$ | TMP Michaelis-Menten<br>constant | 0.52 |
| $h_3$ | Hill coefficient for TMP | 1.05 |

### EF1a-rtTA_TRE3G-mCherry model

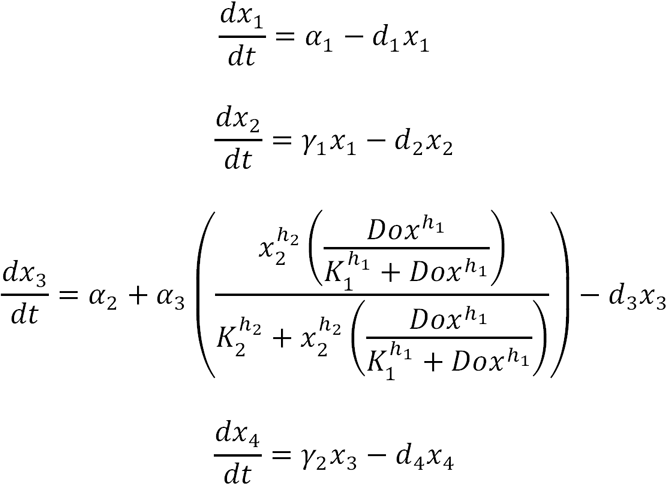

### EF1a-rtTA_TRE3G-DDmCherry model

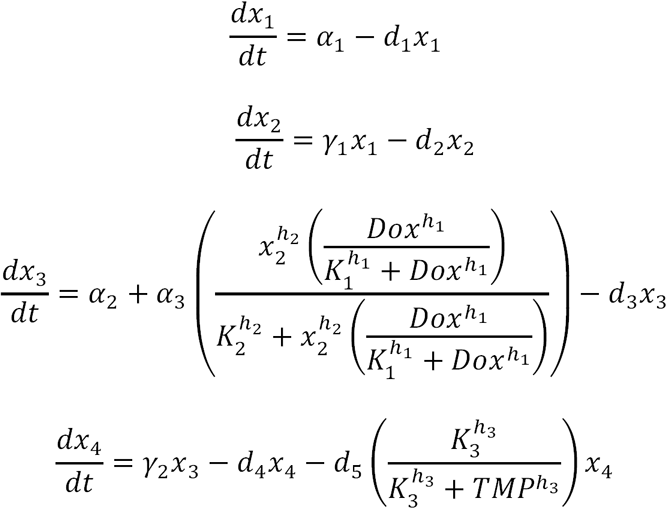

### EF1a-DDrtTA_TRE3G-mCherry model

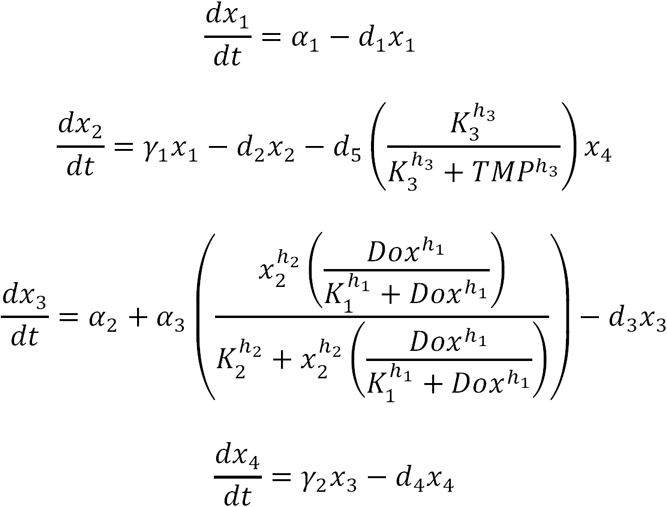

### EF1a-DDrtTA_TRE3G-DDmCherry model

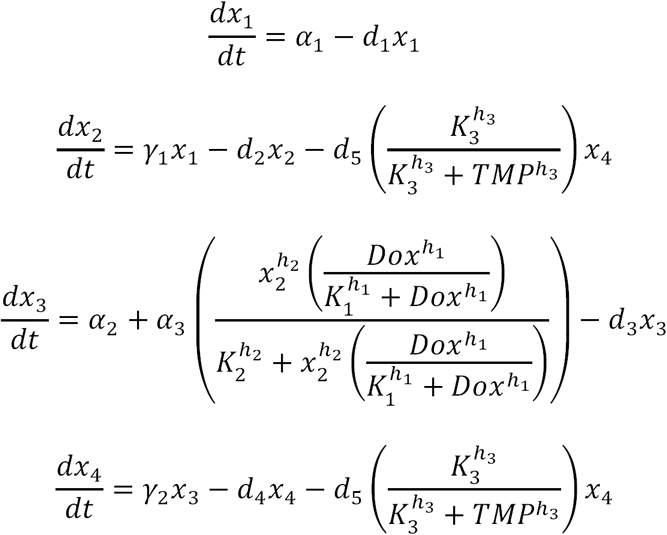

### Microfluidics/microscopy-based time lapses

Microfluidics/microscopy-based experiments were performed using the device designed and optimised for mammalian cells in the laboratory of Prof Jeff Hasty at the University California in San Diego^2^. The topology of the device ensures controlled flow perfusion, minimised cell stress and controlled CO_2_ diffusion. As described in1, cells from a sub-confluent 60cm petri dished were washed with sterile Phosphate-Buffered Saline (PBS, Gibco), trypsinised for 2-3’ at room temperature and centrifuged at 1000rmp for 5’. Pelleted cells were resuspended in 200uL of complete mESC media supplemented with Doxy and TMP and chip-loaded. Before loading, the chip was fulfilled with media containing both inducers from port 5 first and port 1 after. Cell suspension was loaded from port 1 while the vacuum applied to ports 3 and 4. The vacuum allows air to be released from chambers, facilitating cell trapping. Cells were cultured for 24hrs in a tissue culture incubator (5% CO2, 37 °C) under constant perfusion with inducer’s containing media. Media was perfused with a syringe directly connected to port 2 *via* 24-gauge PTFE tubing (Cole-Parmer Inc.). Port 5 was used for waste media whereas ports 1, 6 and 7 were plugged to avoid media spillage. The day after, the device was transferred on the widefield microscope for *in silico* feedback control experiments. The actuation system consists of two motor-controlled syringes (http://biodynamics.ucsd.edu/dialawave/) connected to port 6 and 7. One syringe always contains Doxy/TMP media, whereas the other contains plain media (Figs. 3c, 4e and Supplementary Fig. 4f), or Doxy- (Fig. 3a) or TMP- (Fig. 3b) supplemented media, depending on experimental set-up. Ports 1, 2 and 5 are also connected to static syringes working as waste tanks. If the height of the waste tanks is fixed, the one of the perfusing media ports is automatically adjusted during the experiment to change the input provided to cells. Input administration was measured using a green dye (1uM Atto488 dye from ThermoFisher), added to Doxy/TMP containing syringe.

### Image segmentation

The segmentation algorithm follows the methodology illustrated in^1^. Briefly, a threshold is defined to generate a binary image selecting only pixels belonging to cell edges. Then, by using dilation and filling operators, it derives a binary image (mask) that selects the portion of the original image covered by cells. The mask obtained is applied to the red field image. In order to calculate the average intensity fluorescence of pixels belonging to cells, the background signal measured in a cell-free portion of the chamber is subtracted from the value of mask pixels.

### Fluorescence microscopy

The control platform consists of a Leica DMi8 inverted microscope equipped with the digital camera Andor iXON 897 ultra back-illuminated EMCCD (512x512 16μm pixels,16 bit, 56 fps at full frame), and an environmental control chamber (PeCon) for long-term temperature control and CO2 enrichment. The Adaptive Focus Control (AFC) ensures focus is maintained during the entire time-course experiment. The experimental set-up includes consecutive acquisition in three channels (phase contrast, green and red fluorescence) with a 20X objective every 60mins.

### Relay control

The Relay control strategy can be expressed as follows:

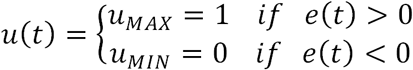

where the control error *e*(*t*) = *r*(*t*) – *y*(*t*) is the difference between the control reference signal *j* and the system output: * is the control input. The Relay controller requires only the computation of the control error at each sampling time *e*(*KT*), where *T =* 60mins, whose sign dictates which input must be provided to the cells. Specifically, cells are treated with inducers-containing medium for the next 60mins if *e*(*KT*) > 0 or medium without inducer otherwise. A 5% hysteresis interval to the controller, corresponding to a tolerance interval around the set-point in which the Relay algorithm ignores the control error, was added to avoid chattering^1^.

In set-point control experiments, we present the response of rtTA_TRE3G-DDmCherry mESCs to three different control inputs, i.e. Doxy (1000ng/ml) in Fig. 3a, TMP (100nM) in Fig. 3b, combination of Doxy (1000ng/ml) and TMP (100nM) in Fig. 3c and Supplementary Fig. 4f, or combination of Doxy (100ng/ml) and TMP (10nM) in Fig. 4e.

This control strategy, although being simple, succeeds in keeping the system output close to the desired reference. Typically, the controlled variable oscillates around the reference and it is acceptable if the oscillation amplitude is sufficiently small^3^.

